# Differential alphavirus defective RNA diversity between intracellular and encapsidated compartments is driven by subgenomic recombination events

**DOI:** 10.1101/2020.03.24.006353

**Authors:** RM Langsjoen, AE Muruato, SR Kunkel, E Jaworski, A Routh

**Author notes:** Address correspondence to Andrew Routh: University of Texas Medical Branch, 6.140A Medical Research Building, 301 University Blvd, Galveston, TX 77555-1061, Lab: (409)772-6680.

## Abstract

Alphaviruses are positive-sense RNA arboviruses that can cause either a chronic arthritis or a potentially lethal encephalitis. Like other RNA viruses, alphaviruses produce truncated, defective genomes featuring large deletions during replication. Defective RNAs (D-RNAs) have primarily been isolated from virions after high-multiplicity of infection passaging. Here, we aimed to characterize both intracellular and packaged viral D-RNA populations during early passage infections under the hypothesis that D-RNAs arise *de novo* intracellularly that may not be packaged and thus have remained undetected. To this end, we generated NGS libraries using RNA derived from passage 1 (P1) stock chikungunya virus (CHIKV) 181/clone 25, intracellular virus, and encapsidated P2 virus and analyzed samples for D-RNA expression, followed by diversity and differential expression analyses. We found that the diversity of D-RNA species is significantly higher for intracellular D-RNA populations than encapsidated and specific populations of D-RNAs are differentially expressed between intracellular and encapsidated compartments. Importantly, these trends were likewise observed in a murine model of CHIKV 15561 infection, as well as *in vitro* studies using related Mayaro, Sindbis, and Aura viruses. Additionally, we identified a novel subtype of subgenomic D-RNA that are conserved across arthritogenic alphaviruses. D-RNAs specific to intracellular populations were defined by recombination events specifically in the subgenomic region, which was confirmed by direct RNA nanopore sequencing of intracellular CHIKV RNAs. Together, these studies show that only a portion of D-RNAs generated intracellularly are packaged and D-RNAs readily arise *de novo* in the absence of transmitted template.

**IMPORTANCE:** Our understanding of viral defective RNAs (D-RNAs), or truncated viral genomes, comes largely from passaging studies in tissue culture under artificial conditions and/or packaged viral RNAs. Here, we show that specific populations of alphavirus D-RNAs arise *de novo* and that they are not packaged into virions, thus imposing a transmission bottleneck and impeding their prior detection. This raises important questions about the roles of D-RNAs, both in nature and in tissue culture, during viral infection and whether their influence is constrained by packaging requirements. Further, during the course of these studies, we found a novel type of alphavirus D-RNA that is enriched intracellularly; dubbed subgenomic D-RNAs (sgD-RNAs), they are defined by deletion boundaries between capsid/E3 and E1/3’UTR regions and are common to chikungunya, Mayaro, Sindbis, and Aura viruses. These sgD-RNAs are enriched intracellularly and do not appear to be selectively packaged, and additionally may exist as subgenome-derived transcripts.

## INTRODUCTION

The *Alphavirus* genus encompasses many important human pathogens, including the equine encephalitis viruses, chikungunya virus (CHIKV), Sindbis virus (SINV), and Mayaro virus (MAYV), among others. Pathogenic alphaviruses are transmitted by a mosquito vector and generally follow the same replication strategy despite infecting distinct organisms and tissues. The alphavirus genome consists of single-stranded positive-sense RNA and encodes 4 non-structural genes, nsP1-4, and 5 structural genes: capsid, E3, E2, TF/6K, and E1. The viral genome is capped and polyadenylated, allowing translation of the non-structural genes upon entry of viral genomic RNA in the cytosol. From this, the viral replication machinery synthesizes anti-sense genomes which are subsequently used to synthesize new genomic sense RNA (1). A key feature of the alphavirus genome is a subgenomic (sg) promoter, which controls expression of sgRNA from anti-sense template RNA independently of its genomic length counterpart. This sgRNA encodes the structural polyprotein, which includes all the structural proteins.

Viral RNA polymerases are notoriously error-prone and with few exceptions lack error-correcting capabilities. While this is most often associated with high mutation rates, these errors frequently include recombination events that cause deletion and/or duplication events (2). This can result in truncated and nonviable genomes that are termed defective-RNAs (D-RNAs) or defective viral genomes (DVGs). D-RNAs are replication defective alone but can be replicated, packaged, and propagated in the presence of wild-type (WT) or helper virus provided they retain the minimally required functional motifs in their genomes (3). Our current understanding of D-RNAs comes largely from *in vitro* passaging studies that use a high multiplicity of infection (MOI, or infectious units/cell) or undiluted virus over the course of multiple viral passages. In this way, D-RNAs have been discovered for nearly every pathogenic RNA virus family in tissue culture (4–12) and are referred to as *defective-interfering* RNAs (or particles) when they have the demonstrated ability to attenuate wild-type virus replication in tissue culture or in animal models of disease. Additionally, they have been proposed to promote viral persistence (13, 14). Although D-RNAs have been considered an epi-phenomenon of cell-culturing practices, emerging deep sequencing technologies have enabled researchers to uncover D-RNAs in both natural and laboratory animal infections (15, 16), as well as in human samples (17, 18).

Through sequencing of *in vivo* D-RNA populations, it is now emerging that D-RNAs play important roles in the outcome of disease in a number of studies. For example, respiratory syncytial virus (RSV) D-RNAs isolated from human nasopharyngeal samples were correlated with heightened immune response to infection and subsequently to improved patient outcomes (17). Similarly, D-RNAs have been isolated from both animals and humans infected with various influenza A virus (IAV) strains (19–22), many of which are similar to those observed *in vitro* (15). Interestingly, a mutation in the viral RNA polymerase that reduced production and subsequent accumulation of D-RNAs was associated with increased influenza disease severity in both humans and mice (19); further, virus isolates from humans with severe IAV infection outcomes contained lower amounts of D-RNAs than those from mild outcomes. Like RSV D-RNAs, IAV D-RNAs are thought to be immunostimulatory and therefore lead to improved patient outcomes (19, 23–25). However, it has also been proposed that high amounts of D-RNAs present in a live-attenuated vaccine formulation can lead to decreased vaccination efficacy by suppressing vaccine strain replication and subsequent immune stimulation (22). Thus, biological functions of D-RNAs, as well as their potential use in vaccines and therapeutics, are of increasing interest.

Alphavirus D-RNAs were first formally described for Semliki Forest virus (SFV) (26, 27) and SINV (28), two prototypical Old World alphaviruses. As with other viral D-RNAs, these alphavirus D-RNAs were generated by undiluted/high MOI serial passaging and were found to suppress wild-type parental virus replication. Later, SINV D-RNA production in persistently-infected BHK cells was positively correlated with increased resistance to SFV challenge (29), indicating a potential for D-RNAs to interfere with the replication cycles of heterologous alphaviruses. The first alphavirus D-RNAs to be sequenced were derived from passage (P) 11 SFV; Lehtovaara *et al* subsequently found that SFV D-RNAs contained conserved nucleotide sequences from the most 5’ and 3’ ends of the genome, which were rearranged across several repeats (30, 31). Similar deletions spanning the majority of the two open reading frames were later identified for SINV (32), although Monroe *et al* additionally found that D-RNA populations were generally heterogeneous and comprised of numerous species. From these and other studies (26–28, 33, 34) it has been hypothesized that alphavirus D-RNAs are encoded by defective negative-sense templates that are subsequently transcribed into defective positive-sense transcripts. This hypothesis is supported by the discovery of novel double-stranded intracellular RNA species in late-passage SINV infections (33), which suggests the presence of a truncated intermediate. Later, a particularly common deletion spanning from the nsP1-E1 genes of the SINV genome was described in the context of a low-fidelity SINV polymerase mutant (35). More recently, CHIKV has been shown to generate recombinant RNAs in tissue culture, especially RNAs featuring complex duplication/deletion events in the 3’UTR (35–40). While D-RNAs have yet been described for CHIKV in either natural or laboratory animal infections, D-RNAs generated *in vivo* have been identified for other alphaviruses. For example, D-RNAs have been found for the distantly related salmonid alphavirus 3 in both salmon from Atlantic farms as well as experimentally infected salmon (16, 41). Additionally, 6K deletion mutants were identified in a Venezuelan equine encephalitis virus (VEEV) isolate from a sentinel hamster (42). Finally, SINV D-RNAs have been recovered from experimentally infected *Drosophila (40)*.

Despite increasing interest in the biological consequences of D-RNA production, the majority of D-RNA research still generally relies on identifying D-RNAs through serial *in vitro* passaging studies. Moreover, all D-RNA studies to date have focused on packaged D-RNA populations, only assaying intracellular compartments after D-RNAs have accumulated over multiple passages. Thus, important questions about the biogenesis and intracellular functions of D-RNAs produced *de novo* remain largely unexplored. In this study, we characterized both intracellular and packaged D-RNA populations during early viral passages, under the hypothesis that numerous D-RNAs arise *de novo* intracellularly that are not packaged. By thoroughly investigating intracellular RNA diversity and its relation to packaged RNA populations, we can define D-RNA subtypes and thereby elucidate specific roles D-RNAs play during infection. To this end, we used Illumina sequencing to sequence RNA from passage (P) 1 alphavirus stock, intracellular RNA (P1.5), and resultant P2 encapsidated RNA, and various bioinformatics approaches to analyze differences in D-RNA diversity and expression. Intracellular D-RNA expression was also evaluated using Oxford Nanopore Technologies’ direct RNA sequencing. Finally, we utilized a murine model of CHIKV infection to demonstrate that these *in vitro* trends hold true in a complex biological system.

## RESULTS

### Intracellular and encapsidated CHIKV D-RNA populations are distinct

To investigate potential differences between intracellular and packaged D-RNA populations during early passages, African green monkey kidney cells (Vero) were infected with CHIKV 181/clone 25 at multiplicity of infection (MOI) of 2. After 12 hours incubation, cells were washed three times with PBS, and then total cellular RNA was extracted and rRNA-depleted. Concomitant infections were incubated for 48 hours, and then supernatant was collected, clarified by centrifugation, and concentrated using PEG-NaCl precipitation; concentrated virus was incubated with 2μg RNase A for 1 hour at room temperature to remove non-packaged RNAs prior to RNA extraction. By following this study design, we were able to maximize available viral RNA from both compartments and thus enable us to identify rare events. RNAseq libraries were then constructed using the previously described ClickSeq library preparation method optimized for the discovery of rare recombination events (43, 44) and sequenced on an Illumina NextSeq550. Reads were trimmed and filtered using *fastp* (45) and analyzed using the *ViReMa* v1.5 pipeline, which provides alignment data, recombination events, and associated count data (20, 44, 46). This study design was repeated once with one replicate and a second time with 3 replicates (used for statistical analyses). Additionally, RNA from two separate stock vials was purified and sequenced. These were not included in statistical comparisons, but are shown none-the-less to establish a baseline D-RNA phenotype. All reactions resulted in robust sequencing read coverage across the reference virus genome, with median nucleotide coverage ranging between 34268 and 71485 reads (Table 1). Median coverage was slightly higher for nucleotides in the subgenomic region in intracellular samples, ranging between 5.4-5.9x higher than those in the nonstructural gene cassette. Overall, after normalizing count data to the number of mapped reads per million sequenced reads, both the total number of recombination events, as well as the number of unique events, were significantly higher among intracellular RNAs than encapsidated RNAs (Student’s T-test, p<0.01 for all; Fig. 1A-B), indicating that the majority of D-RNAs generated during replication were not packaged. This is supported by Shannon’s entropy analysis, calculated from raw D-RNA count data weighted against median wild-type coverage as previously described (44), which found that Shannon’s diversity index (*H*) for D-RNAs is significantly higher for intracellular RNAs than encapsidated (Student’s T-test, p<<0.005; Fig. 1C); thus, intracellular CHIKV D-RNA populations are more diverse than encapsidated, indicating a bottleneck imposed by the packaging process. The distribution of recombination event types (i.e., those with donor sites in the nonstructural/genomic coding region, in the structural/subgenomic coding region, and within the 3’UTR) also differs between intracellular and encapsidated compartments, with 3’UTR events generally enriched in the encapsidated compartment and subgenomic events highly enriched in the intracellular compartment (Fig. 1D).

**Table 1.**
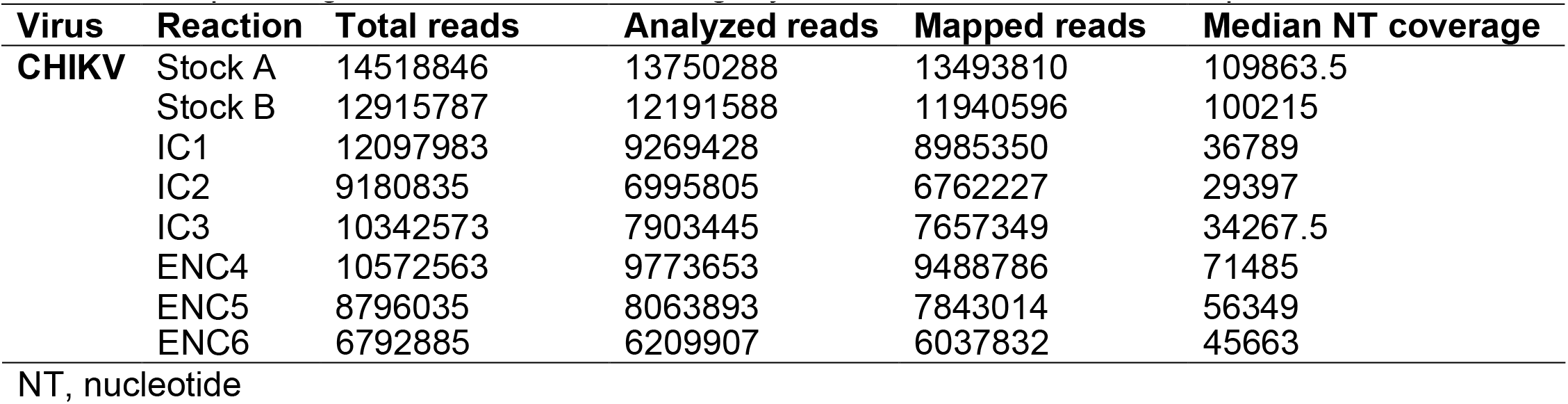
Sequencing reaction data for chikungunya virus 181/clone25 ClickSeq libraries

**Figure 1.**
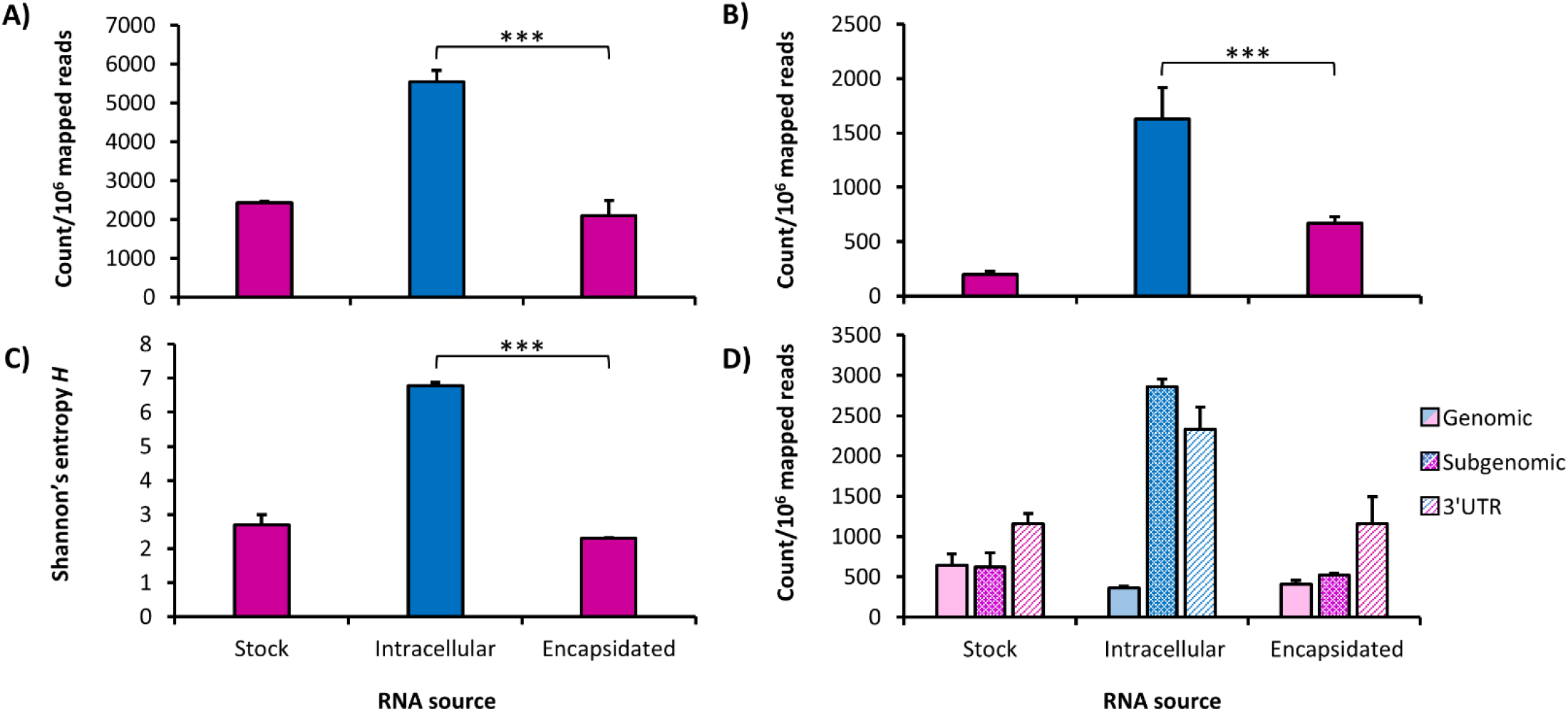
Overview of D-RNA expression in P1 stock, intracellular, and P2 RNA samples. RNA from two replicates of Passage 1 (P1) stock virus, three replicates of intracellular (P1.5), and three replicates of resulting Passage 2 (P2) encapsidated RNA from CHIKV-infected Vero cells and supernatant were sequenced and analyzed for D-RNA expression using *ViReMa v1.5*. D-RNA count data were then normalized to count per 10^6^ reads and **(A)** total number of D-RNAs/10^6^ mapped reads **(B)** unique D-RNAs/10^6^ mapped reads calculated for each compartment. This was followed by calculating **(C)** Shannon’s diversity index *H*, weighted for median nucleotide coverage, using a custom python script. Pink indicates samples taken from encapsidated virus (P1 stock and P2), blue indicates intracellular samples. One asterisk represents p<0.05, two asterisks represents p<0.01, three asterisks represents p<0.001 (Student’s T-Test). **D)** The types of total recombination events were further broken down into genomic (those with donor sites in the nonstructural coding region), subgenomic (those with donor sites in the structural coding region), and 3’UTR (those with donor and acceptor sites within the 3’UTR) for each RNA source.

A principle component analysis (PCA) of recombination event count frequencies was performed for intracellular and encapsidated D-RNAs (Fig. 2A), which revealed that intracellular samples cluster together while two of the three encapsidated replicates cluster together with the third clustering closer to intracellular samples, indicating that variation in D-RNA expression patterns is mostly unique to its respective sources (intracellular vs. encapsidated). This is supported by differential expression analysis (Fig. 2B and C), performed with DESeq2 (47) followed by hierarchical clustering with Cluster 3.0 (48) and TreeView (49), which revealed that D-RNAs with recombination events found in distinct genomic regions are differentially enriched in each RNA source. Of note, recombination events enriched intracellularly occur within the subgenomic region. Altogether, these data demonstrate that intracellular and encapsidated D-RNA populations differ significantly by all metrics employed, confirming that many D-RNAs arise *de novo* that are not efficiently packaged.

**Figure 2.**
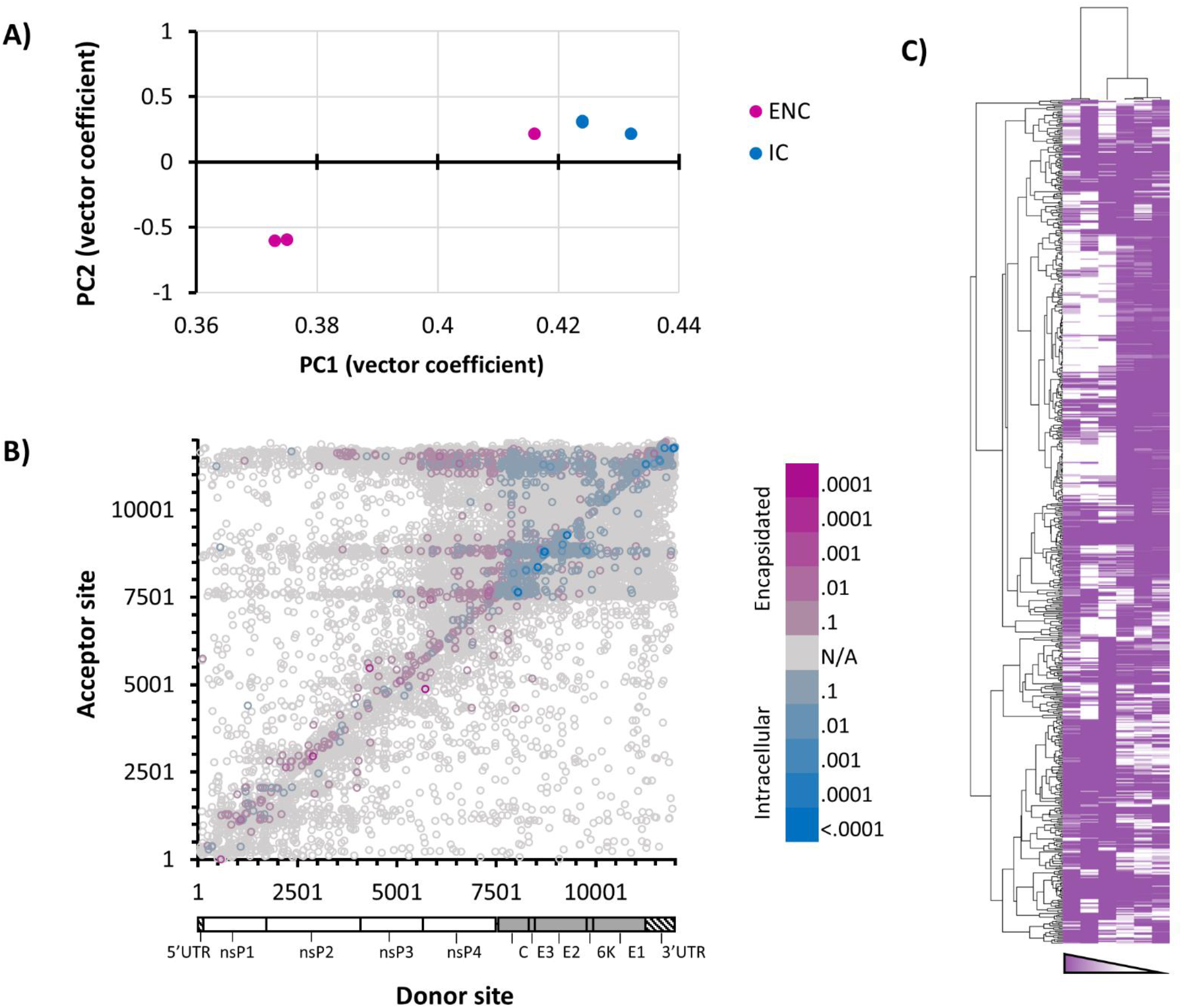
Characterization of CHIKV D-RNA populations using differential expression approaches. Three replicates each of intracellular (blue) and resulting encapsidated (pink) RNA from CHIKV infected Vero cells and supernatant, respectively, was sequenced and evaluated for defective RNA (D-RNA) expression using *ViReMa* v1.5. Count tables of D-RNAs abundance were passed to DESeq2 for subsequent analyses. **A)** Two-dimensional principle component analysis **B)** differential expression of all D-RNA species **C)** hierarchical clustering of CHIKV D-RNA species (using DESeq2 output with Cluster 3.0 and Treeview)

### Subgenomic deletions: a new D-RNA archetype for CHIKV

To compare specific boundaries, all normalized ViReMa count data from each data set (stock, intracellular, and encapsidated) were combined; normalized count data for D-RNAs present in at least two out of three samples were averaged, and then the boundaries for D-RNAs with greater than 2 counts per 10^6^ mapped reads are illustrated using heatmaps, along with associated coverage data (Fig 3). While several distinct and abundant acceptor sites (indicated by horizontal striations) appear in all three data sets around nucleotides 7750, 8850, and 11500 (Fig. 4B), subgenomic recombination events are enriched among intracellular RNAs compared to both stock and encapsidated samples. To further visualize this, the top 80 recombination events for intracellular and encapsidated RNAs were divided into “duplication”/“back-splicing” events and deletion events and then mapped (Fig. 5, one representative replicate shown). Several major D-RNA archetypes emerge: deletion events between nsP4 region and the end of E1/3’UTR, which are common among both intracellular and encapsidated RNAs; deletion events between capsid-E3 and E1-3’UTR, which are common intracellularly but are relatively rare among encapsidated RNAs; and deletion and duplication events occurring specifically within the 3’UTR, which are among the most common in both intracellular and encapsidated RNAs (subgenomic recombination types among intracellular RNAs are illustrated in Fig. S1). This illustrates that the composition of intracellular and encapsidated D-RNA populations differs substantially, while also demonstrating novel D-RNA archetypes not previously observed in alphaviruses.

**Figure 3.**
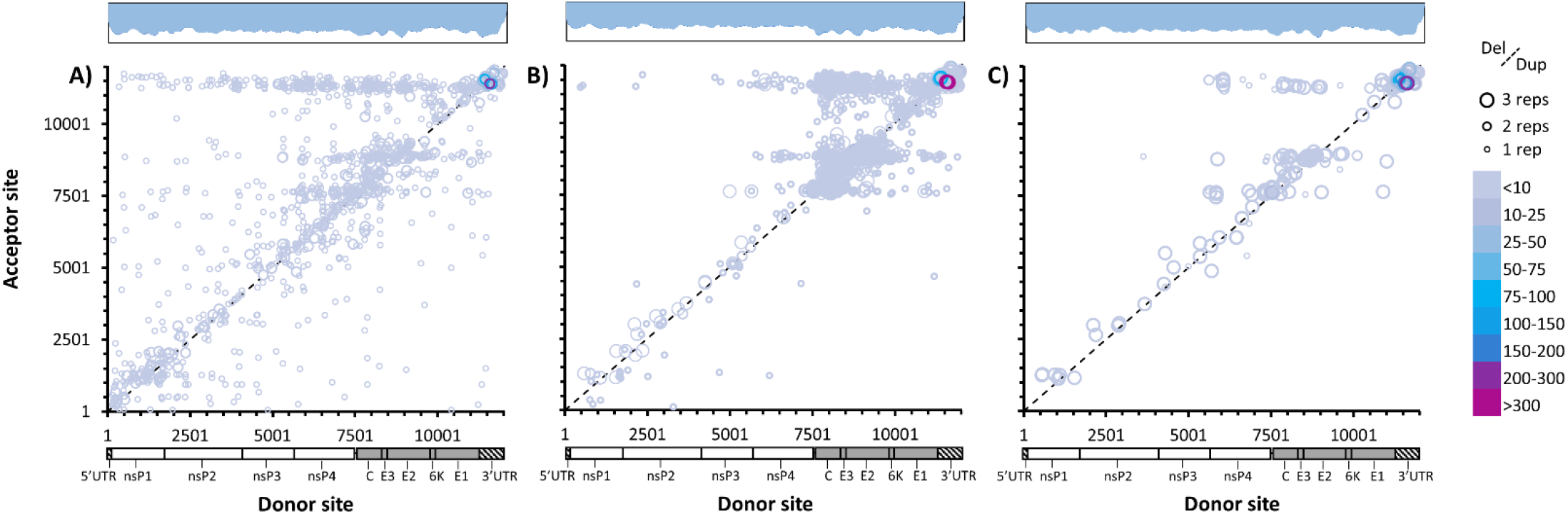
Recombination boundaries observed among CHIKV D-RNAs. CHIKV recombination junctions for D-RNAs with ≥1 count per every 2×10^6^ reads. RNA was derived from **A)** P1 stock virus (two replicates) **B)** intracellular RNA (three replicates) from infected Vero cells and **C)** resulting P2 encapsidated virus (three replicates). Circle size indicates number of replications, line delineates deletion vs. duplication/insertion events, while color indicates count/10^6 mapped reads. Log coverage data shown above, with light blue indicating average coverage over three replicates and dark blue the standard deviation.

**Figure 4.**
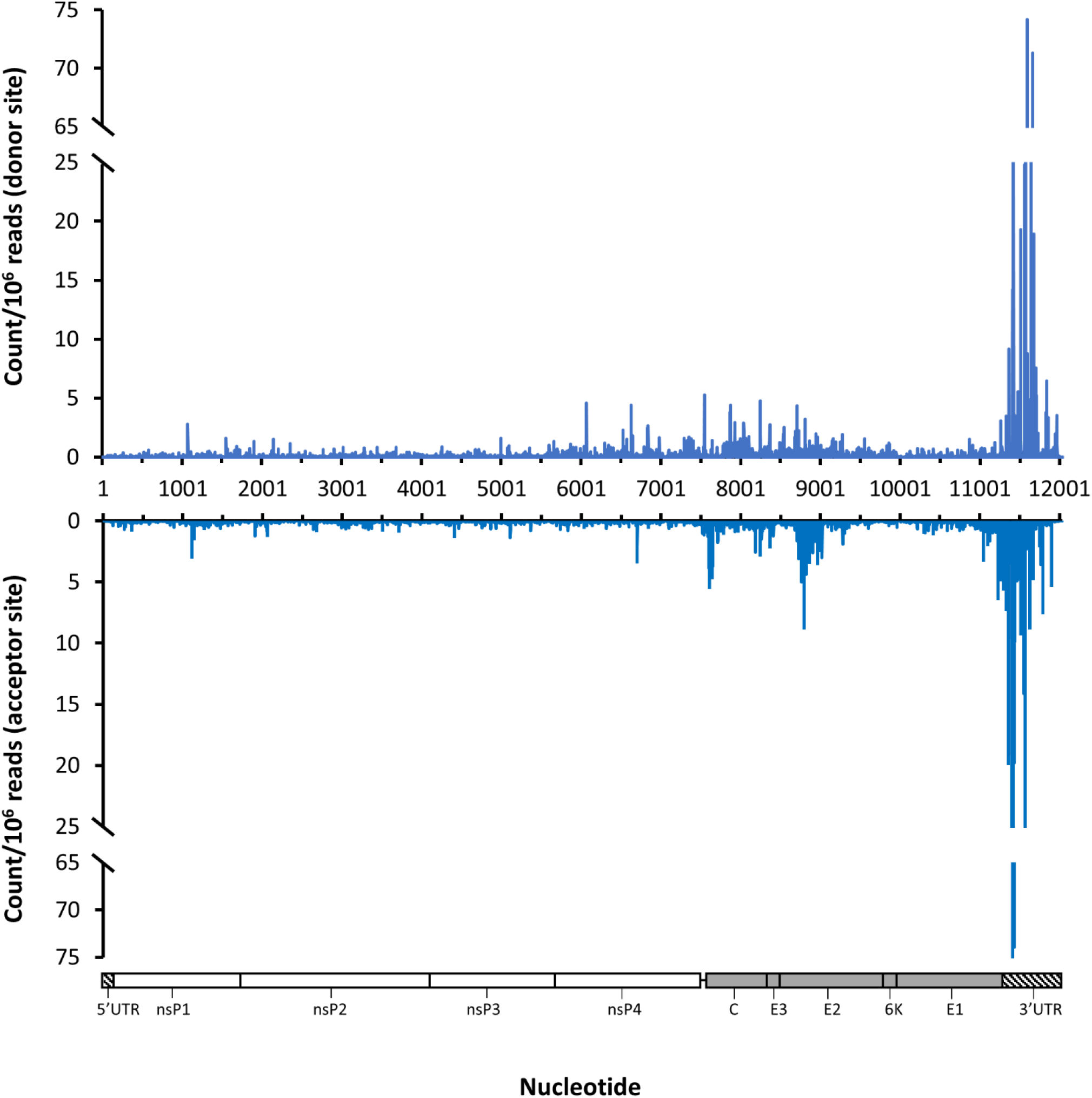
Recombination junction position counts for CHIKV D-RNAs. Three replicates of intracellular RNA from chikungunya virus (CHIKV)-infected Vero cells was collected 12 hours post-infection and then sequenced and evaluated for defective RNA expression using *ViReMa* v1.5. Recombinant count data were normalized to count per 10^6^ CHIKV-mapped reads, and then average nucleotide count for donor sites **(Top)** and acceptor sites **(bottom)** were plotted to identify recombination hotspots

**Figure 5.**
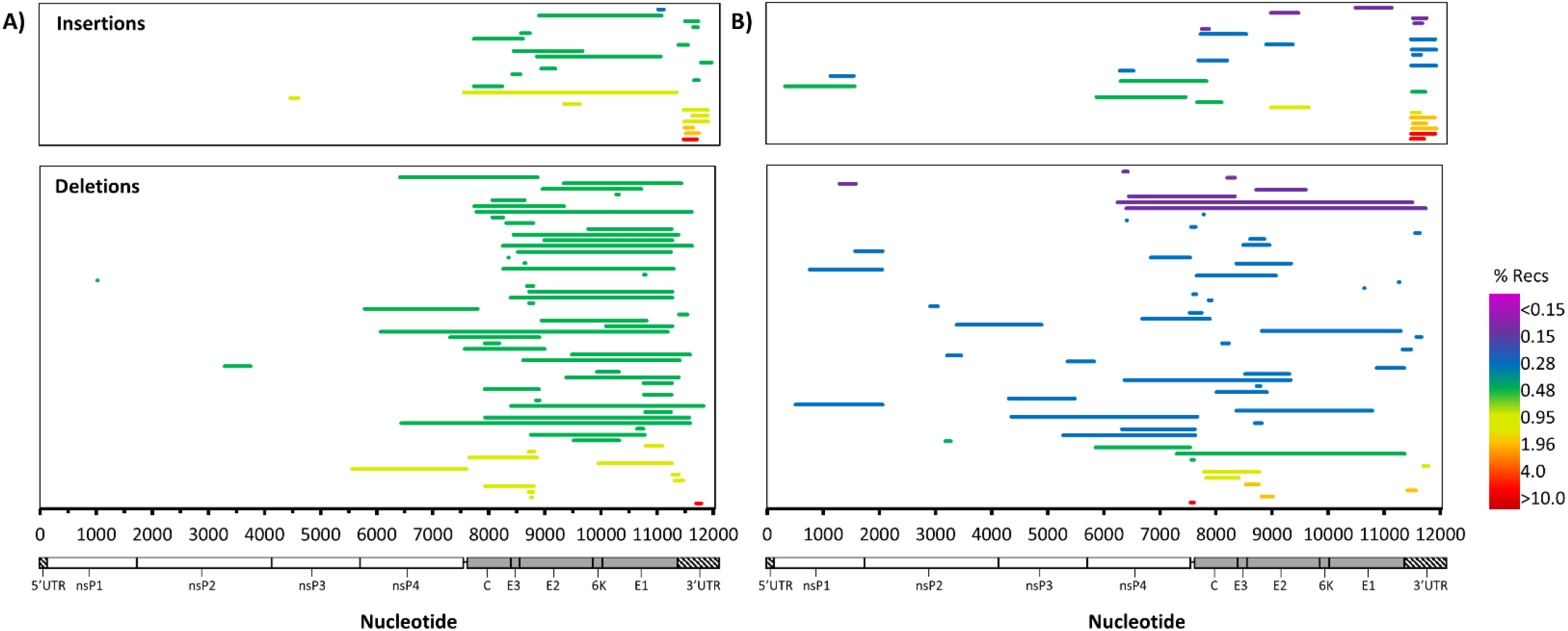
Top recombination events observed among CHIKV 181/clone25 D-RNAs. One replicate of each of **(A)** intracellular RNA from CHIKV infected Vero cells collected 12 hours post-infection (HPI) and **(B)** supernatant collected 48 HPI, respectively, was sequenced and evaluated for defective RNA (D-RNA) expression using *ViReMa* v1.5. Recombinant count data were normalized to count per 10^6^ CHIKV-mapped reads and the top 80 recombination events were split into insertion/duplication events (top) and deletion events (bottom). Color represents the percentage of overall recombination events represented by a single event.

### Direct RNA sequencing of CHIKV RNA differentiates subgenomic D-RNAs and genomic D-RNAs

For the recombination events observed in the subgenomic region, it is possible that they occur on either genomic or sg transcripts intracellularly; additionally, it is difficult to quantify relative numbers of D-RNAs to full-length transcripts using short-read sequencing due to, for example, variance inherent to RNAseq in nucleotide coverage across the virus genome and at boundaries of recombination events. Thus, to put these sgDels into context, two separate Oxford Nanopore Technologies’ (ONT) Direct RNA Sequencing (DRS) libraries were generated using two intracellular samples derived from independent infections that utilized completely distinct CHIKV 181/clone25 stocks. ONT DRS directly sequences the input RNA on a single molecule by molecule basis (there is no PCR or amplification process), maintaining native features of the RNA (50). Therefore, this technology can robustly quantify relative numbers of genomic and sgRNA reads. Reads were based-called using Albacore v2.0.1 and aligned to the CHIKV 181/clone 25 genome using minimap2 (51). Full-length reads were recovered (totaling 75-82 reads per data set), and median nucleotide coverage ranged from 229-249.5 with decreasing coverage from the 3’UTR (due to 3’ to 5’ sequencing from the poly(A) tail) and a sharp drop in coverage following the 5’UTR of the sgRNA (Fig. S2), showing a clear difference between genomic and subgenomic reads. From aligned data, specific read junctions were extracted from the SAM file and reads containing deletions were assessed (Table 2). For both sequencing data sets, multiple deletion events on a single read were rare, with 96-98% of deleted reads featuring just one deletion.

**Table 2.**
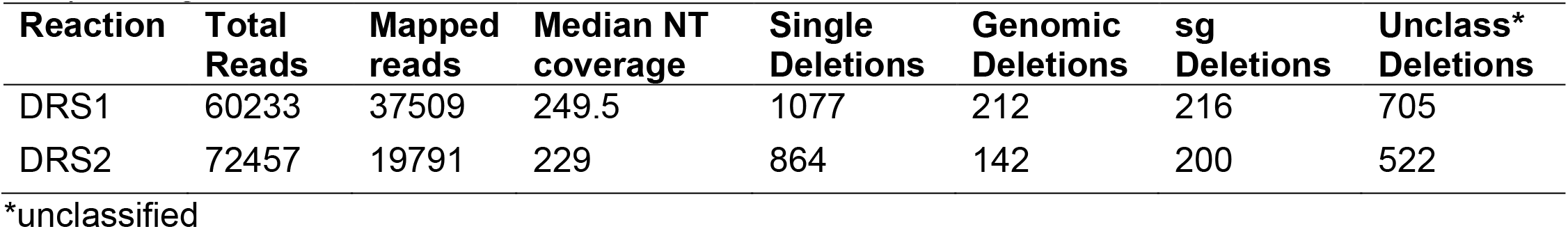
Sequencing reaction data for chikungunya virus 181/clone25 direct RNA sequencing libraries

First, reads were placed into one of two categories: 1) genomic reads containing deletions, defined as a read beginning between NT 1-7450; and 2) sg reads containing deletions, defined as a read beginning between NT 7500-7550. Beyond these nucleotides, it is not possible to discern between true sgRNA reads and genomic reads that may have simply been truncated during the sequencing process (either due to RNA fragmentation or incomplete 3’ to 5’ nanopore sequencing of the RNA). The largest deletions for genomic and sgRNA reads are shown for one of the libraries (Fig. 6, one representative replicate shown), along with read/deletion boundary counts. These data confirm that many of the sgDels occur specifically on sgRNAs rather than genomic RNAs. Interestingly, although sgDels were present in both genomic and sgRNA transcripts, there was little to no commonality between sgDels observed in genomic reads and sgDels observed in sg reads, with only two sgDels common to both sgRNA and genomic RNA reads in DRS1. Thus, we were unable to confirm the presence of a common defective template for the overwhelming majority of sgDels observed.

Finally, these data additionally reveal that 2.9-3.2% of all aligned reads from intracellular samples contained at least one deletion, calculated by dividing [mapped reads with deletions]/[total mapped reads]. Using this same calculation with Illumina data, the overall percentage of reads containing any recombination event from intracellular samples ranged from 0.2642-0.2842%. This difference is likely due to reverse transcription- and PCR-based biases associated with the Illumina platform (52–54), which can cause variation in nucleotide coverage across the genome and thus necessitate the consideration of local nucleotide coverage when calculating percent D-RNAs. Calculating percent D-RNAs for alphaviruses is further complicated by the molar disparity between genomic and sgRNA expression. Adjusting Illumina count percentages by average nucleotide coverage for a specific region—genomic or subgenomic—suggests that D-RNAs may comprise 9.79±1.25% of intracellular populations and 7.65±1.26% of encapsidated populations. However, there is no universally accepted method to calculate D-RNA proportions using short-read data whereas proportions can be calculated from DRS data using straight-forward ratios (55).

**Figure 6.**
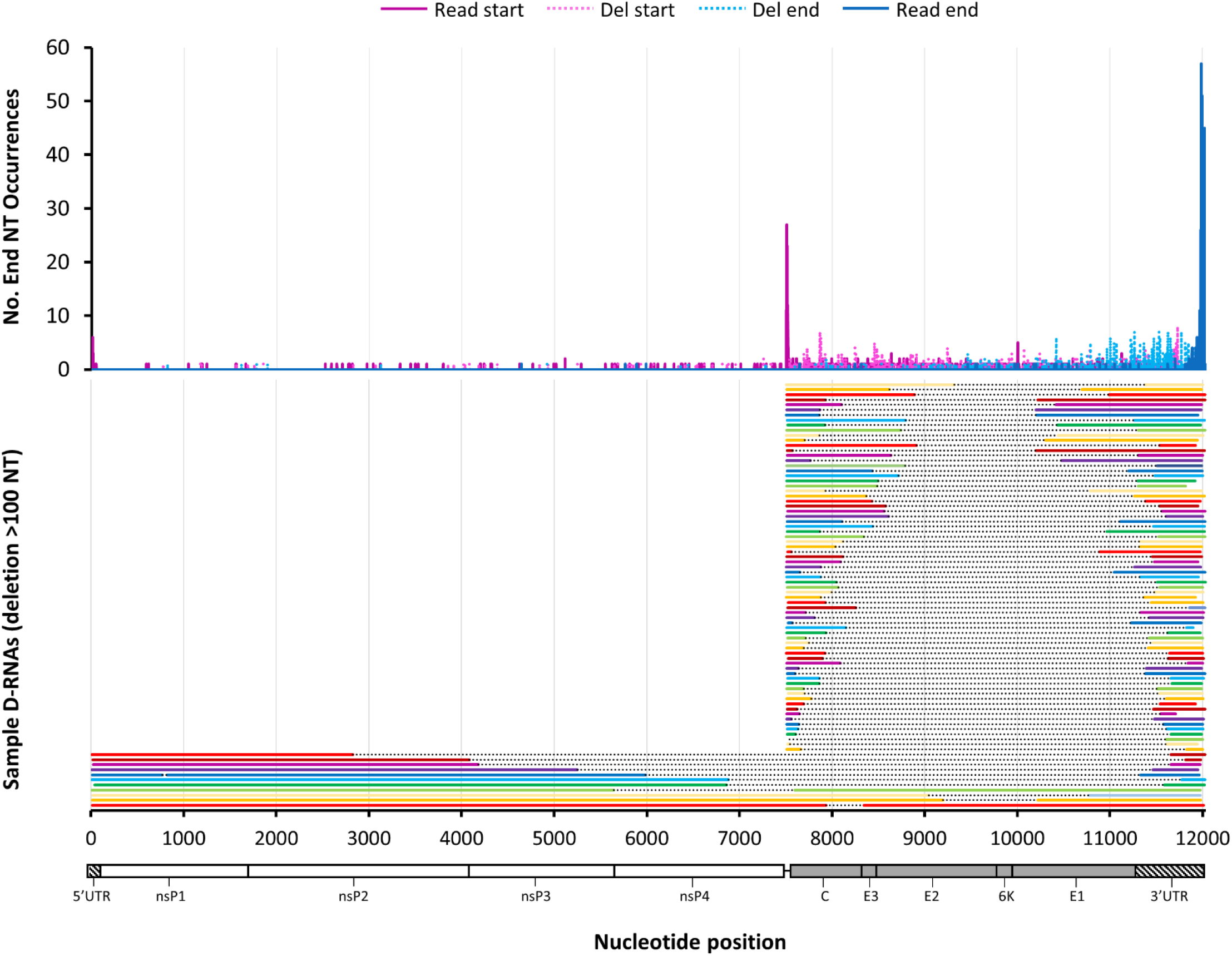
Oxford Nanopore’s direct RNA sequencing of intracellular CHIKV D-RNA species. Intracellular RNA from CHIKV-infected Vero cells was sequenced using Oxford Nanopore’s Direct RNA sequencing kit on a MinION sequencer, reads were mapped using minimap2, and then read boundaries for CHIKV-mapped reads were extracted from SAM files. Boundary counts for defective RNA reads containing deletions were enumerated and plotted **(top)** while the raw single reads containing the largest deletions for genomic and subgenomic D-RNAs are shown **(bottom)**.

### D-RNA expression patterns are conserved across the alphavirus family

To ascertain whether sgD-RNAs are a CHIKV-specific phenomenon or conserved broadly amongst arthritogenic alphaviruses, the above *in vitro* studies were repeated using Mayaro virus (MAYV), Sindbis virus (SINV), and Aura virus (AURV; Table 3). Of these, MAYV shares the highest sequence identity with CHIKV at 62.91% and subsequently groups in the same Semliki Forest virus complex, while SINV and AURV group phylogenetically with the adjacent Western equine encephalitis virus (WEEV) complex (56). Additionally, AURV is the only alphavirus that has been shown to package its sgRNA to date (57–59).

**Table 3.**
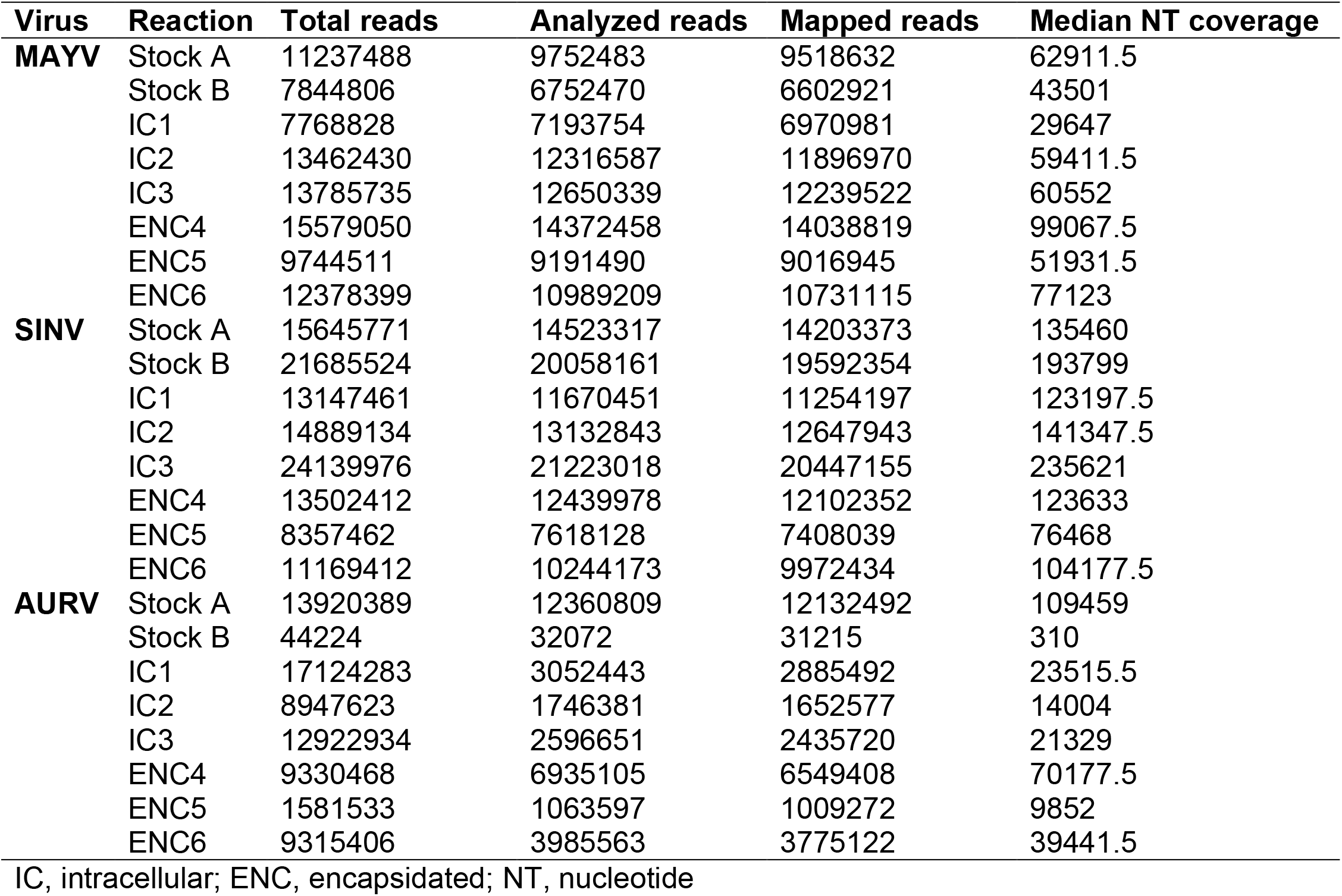
Sequencing reaction data for alphavirus ClickSeq libraries

For all three additional alphavirus species, D-RNA diversity was significantly higher intracellularly than among encapsidated RNAs (table 4), again indicating both a packaging bottleneck in the transmission of D-RNAs, as well as the potential for D-RNAs to arise *de novo* without the aid of a transmitted template. Interestingly, the two New World representatives, MAYV and AURV, shared similar intracellular and encapsidated D-RNA diversity indices, whereas the two Old World representatives, CHIKV and SINV, shared similar intracellular, but not encapsidated, indices. Further, these differences in population diversity, as with CHIKV, were largely driven by recombination events in the subgenomic region for MAYV (Fig 7). Recombination events in the SINV subgenome were similarly enriched in intracellular samples, while an additional number of recombination events in and across the nonstructural cassette were significantly enriched in encapsidated samples, including previously described D-RNAs (35). AURV appeared to have a bimodal distribution of differentially expressed D-RNAs, thus making interpretation of these results unreliable. PCA indicated that intracellular and encapsidated samples clustered independently for both MAYV and SINV. However, while encapsidated AURV samples clustered together, intracellular samples showed no clustering pattern.

**Table 4.** Shannon’s diversity index *H* for alphavirus infections in vero cells

| Virus | Intracellular *H* | Encapsidated *H* | t | p |
| --- | --- | --- | --- | --- |
| Chikungunya | 6.78±0.1 | 2.31±0.01 | -79.56 | <<0.0001* |
| Mayaro | 5.35±0.07 | 3.26±0.67 | -5.386 | 0.0067* |
| Sindbis | 8.09±1.59 | 3.97±0.02 | -4.503 | 0.0108* |
| Aura | 5.17±0.011 | 3.03±0.42 | -8.518 | 0.001* |

Numbers shown are average ± standard deviation. Test values t and p derived from Student’s T-test.

**Table 5.** Sequencing reaction data for chikungunya virus AF15561 ClickSeq libraries

| Organ | # | Total reads | Analyzed reads | Mapped reads | Median NT coverage |
| --- | --- | --- | --- | --- | --- |
| Serum | 1 | 1991095 | 372210 | 361998 | 2997.5 |
|  | 2 | 4914494 | 3914465 | 3777153 | 31977.5 |
|  | 3 | 10150263 | 8628164 | 8370889 | 71524 |
|  | 4 | 7835883 | 6761112 | 6514128 | 55618 |
| SKM-CL | 1 | 12920938 | 44073 | 42448 | 294 |
|  | 2 | 10884345 | 1538267 | 1481545 | 12460 |
|  | 3 | 12120048 | 236946 | 228495 | 1867.5 |
|  | 4 | 12243525 | 309850 | 298029 | 1863 |
| **SKM-IS** | 1 | 12231654 | 2893892 | 3787566 | 20349 |
|  | 2 | 15379613 | 1665210 | 1593002 | 14151 |
|  | 3 | 14997585 | 1297009 | 1242956 | 10213 |
|  | 4 | 8801363 | 1739616 | 1668388 | 13525 |
| **Heart** | 1 | 13286951 | 225684 | 219139 | 1182 |
|  | 2 | 11620802 | 273366 | 265887 | 1459.5 |
|  | 3 | 18709794 | 223552 | 223552 | 1388 |
|  | 4 | 8152377 | 37435 | 35990 | 216 |
| **Kidney** | 1 | 13810118 | 277261 | 267856 | 1553 |
|  | 2 | 11063493 | 54343 | 52743 | 267 |
|  | 3 | 16439432 | 297215 | 297215 | 2017 |
|  | 4 | 8037635 | 31460 | 30165 | 198 |
| **Spleen** | 1 | 6362763 | 9324 | 8878 | 75 |
|  | 2 | 10006661 | 1548 | 1508 | 0 |
|  | 3 | 8877156 | 730 | 655 | 0 |
|  | 4 | 1009068 | 148 | 147 | 0 |
| **Liver** | 1 | 16473789 | 19659 | 19007 | 141 |
|  | 2 | 15456139 | 13516 | 13004 | 88 |
|  | 3 | 13565351 | 9170 | 8936 | 64 |
|  | 4 | 1652843 | 484 | 468 | 0 |

NT, nucleotide

**Figure 7.**
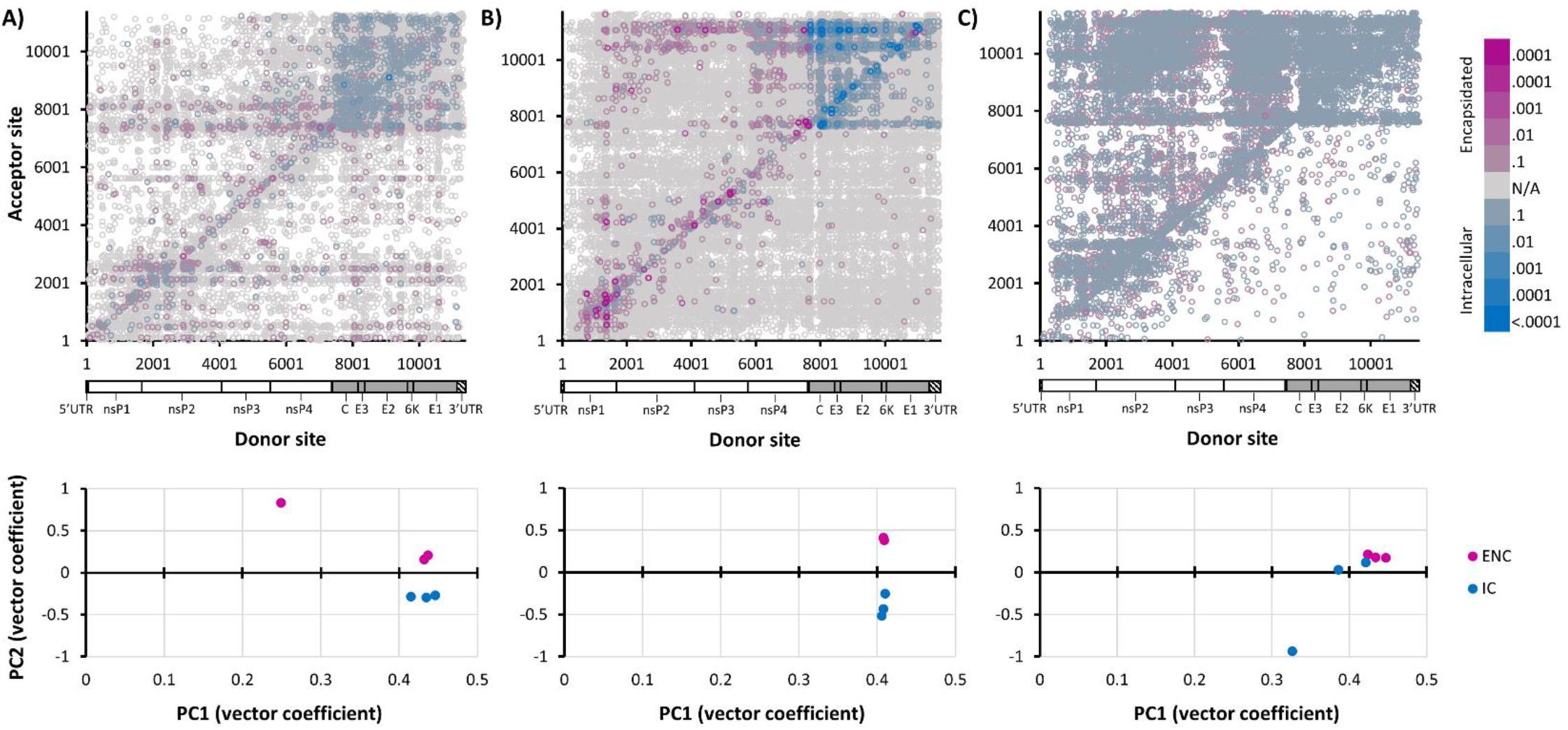
Characterization of alphavirus D-RNA populations using differential expression approaches. Three replicates each of intracellular (blue) and resulting encapsidated (pink) RNA from Mayaro virus **(A)**, Sindbis virus **(B)**, and Aura virus **(C)** infected cells and supernatant, respectively, was sequenced and evaluated for defective RNA (D-RNA) expression using *ViReMa* v1.5. DESeq2 was then used to concatenate and normalize count data for subsequent analyses. Differential expression of all D-RNA species are shown in top panels, while principle component analyses are shown in bottom panels.

Furthermore, heatmaps of recombination junctions reveal that each of these additional alphaviruses showed a strong junction 2 near the subgenomic promoter that carried over from stock through encapsidated RNAs (horizontal striations in Fig. 8), similar to that observed for CHIKV: for MAYV, near NT 7600 (Fig. S3); for SINV, near NT 7760 (Fig. S4); and for AURV, near NT 7840 (Fig. S5). These sites were accompanied by an additional downstream donor site that was nearly absent in both stock and encapsidated RNAs, recapitulating the pattern observed for CHIKV in both *in vitro* and *in vivo* samples. However, while MAYV especially exemplifies this pattern, SINV showed signs of a donor site in both stock and encapsidated samples. MAYV showed several additional acceptor sites in the non-structural genes, as well as one acceptor site near NT 8030. Unlike all other alphaviruses, an acceptor site was observed near the end of the MAYV E1 gene in intracellular samples and only weakly in stock virus. On the other hand, SINV and AURV both displayed acceptor sites near the end of their respective E1 genes in all three sample types. AURV additionally displayed a variety of both donor and acceptor sites and heavily favored deletions over “duplications” in all sample types, aggregating in hotspots in the nsP2 and nsP4 genes.

**Figure 8.**
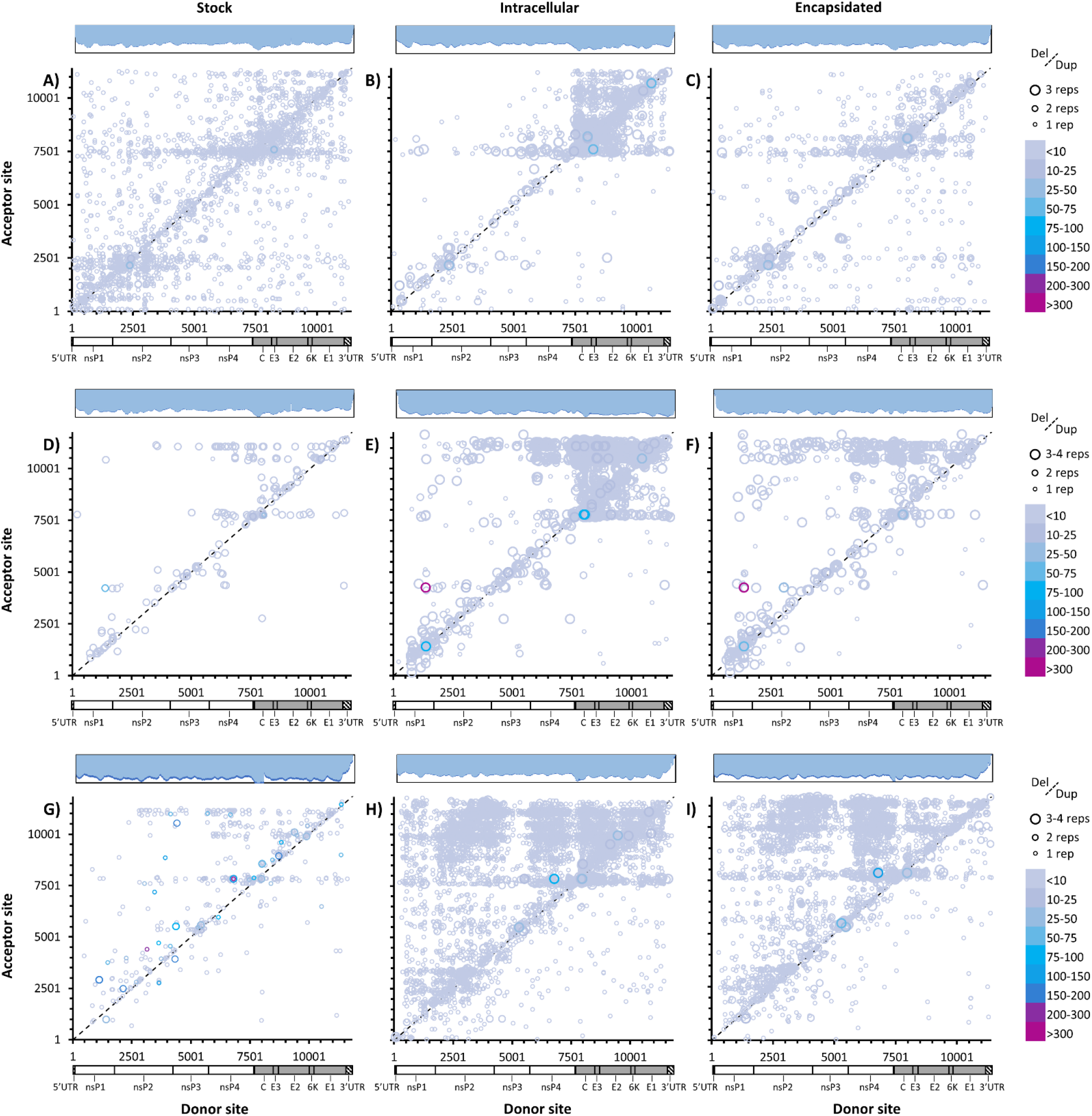
Recombination boundaries observed among alphavirus D-RNAs. Alphavirus recombination junctions for D-RNAs with ≥1 count/2×10^6 reads from stock (left panels), intracellular (center panels), and encapsidated (right panels) for the following viruses: Mayaro virus **(A-C)**, Sindbis virus **(D-F)**, and Aura virus **(G-I)**. Circle size indicates number of replications, line delineates deletion vs. duplication/copyback events, while color indicates count/10^6 mapped reads. Log coverage data shown above, with light blue indicating average coverage over three replicates and dark blue the standard deviation.

### Murine infection with wild-type CHIKV results in similar D-RNA expression patterns as early-passage vero infections

To assess whether *in vitro* D-RNA expression patterns are recapitulated *in vivo*, IFNαR^-/-^ (A129) mice were infected with 10^3^ plaque forming units (PFU) CHIKV 15561, the parental strain to CHIKV 181/clone 25 (60, 61), and monitored for 4 days post-infection (DPI) until clinical signs such as ruffled fur and hunched posture appeared; serum, skeletal muscle from injected and contralateral limbs, heart, kidney, liver, and spleen were collected upon humane euthanization at 4 DPI. Virus in serum was quantified by plaque assay, while RNA was extracted from all samples using either TRIzol or Qiagen RNeasy Fibrous Tissue kit. Ribosomal RNA was depleted from tissue samples and then ClickSeq libraries synthesized for NGS. Fewer than 10,000 reads from most liver and spleen samples mapped to the CHIKV genome despite robust sequencing reactions (Table 4); therefore, spleen and liver were excluded from downstream analyses.

D-RNAs were identified in all tissues as well as serum, featuring the same recombination events as identified during *in vitro* studies. Consistent with our *in vitro* results, D-RNA diversity (as indicated by Shannon’s entropy) was higher in tissues (i.e. intracellularly) than in serum (encapsidated RNAs; Fig. 9A). Further, average serum diversity and average cell-derived encapsidated diversity indices were similar at 1.92 and 2.3, respectively. However, average tissue D-RNA diversity, ranging between 3.07-3.4, was lower than average cell-derived intracellular diversity at 6.8. While only heart muscle and skeletal muscle at the injection site had significantly higher diversity than serum (Kruskal-Wallis with Dunnet’s post-hoc, p<0.05), these animals were not perfused prior to tissue collection and thus excess serum virus may have diluted the diversity of some tissue samples and subsequently affected standard deviation. Murine samples in general failed to cluster by organ (Fig. 9B) with the exception of heart samples, after PCA. Most tissue samples formed a single large cluster with one of the serum samples (from mouse 2), although serum samples in general showed no clustering pattern at all. Thus, although statistical significance was not achieved in all cases, expression trends generally agree with those observed *in vitro*.

**Figure 9.**
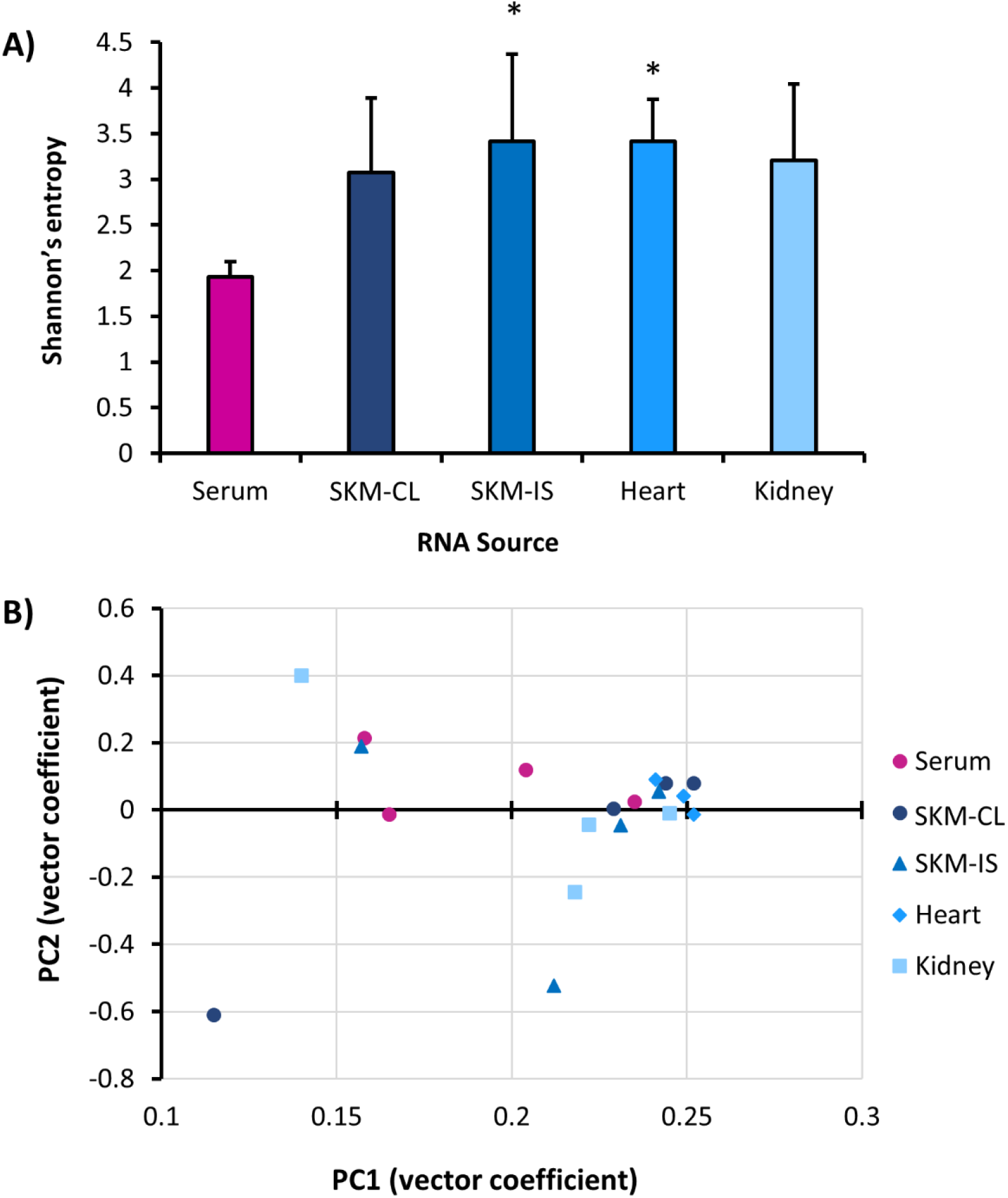
CHIKV D-RNA diversity and expression in a mouse model of infection. Four A129 mice were infected with CHIKV AF15561, and then serum and organs collected 4 days post-infection. RNA was then extracted from RNA and tissues, and then sequenced on an Illumina NextSeq550. Defective-RNAs (D-RNAs) were identified and quantified using *ViReMa v1.5*. From raw data, Shannon’s diversity index *H* **(A)**, weighted for median nucleotide coverage, was calculated for individual data sets using a custom python script (Kruskal-Wallis with Dunnet’s post-hoc, one asterisk indicates p<0.05). Additionally, D-RNA count data analysed using DESeq2, followed by principle component analysis **(B)**.

Also consistent with *in vitro* results, heat maps of recombination junctions reveal the same three acceptor sites observed *in vitro* across both serum and tissue samples, as well as a similar donor site specific to tissues rather than serum (Fig. 10). Interestingly, skeletal muscle, and especially kidney tissue, was particularly enriched for sgD-RNAs. 3’UTR recombination events remained the most common type of D-RNA in all samples. This is also evident when comparing the top deletion events across different tissues, where kidney samples unsurprisingly most closely recapitulate *in vitro* data (Fig. S2-3). In addition to the 3’ boundary observed in the 3’UTR, all mouse samples showed an additional 5’ boundary in the 3’UTR. Not only did we observe similar trends in diversity measures between *in vitro* and *in vivo* studies, but we also observed similar trends in D-RNA population composition. Thus, we have demonstrated biologically relevant D-RNA phenomena that can be exploited for better understanding RNA virus replication with potential applications in human health.

**Figure 10.**
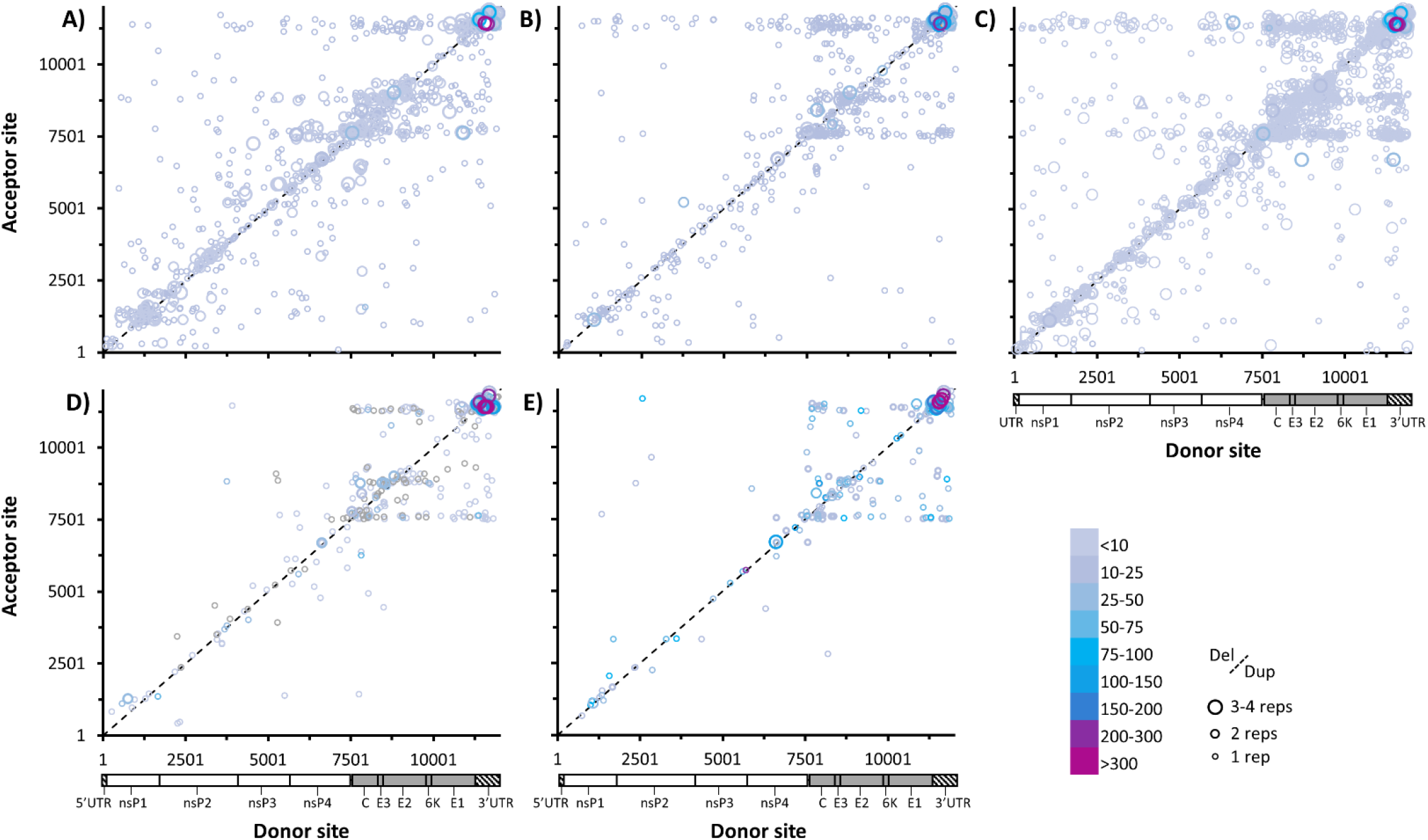
Recombination boundaries observed among CHIKV D-RNAs in a murine model of infection. Four A129 mice were infected with CHIKV AF15561, and then serum and organs collected 4 days post-infection. RNA was then extracted from RNA and tissues, and then sequenced on an Illumina NextSeq550. Defective-RNAs were identified and quantified using *ViReMa v1.5*. CHIKV recombination junctions for D-RNAs with ≥1 count per every 2×10^6^ reads were then plotted for **A)** serum **B)** contralateral skeletal muscle **C)** injection site skeletal muscle **D)** heart and **E)** kidney. Circle size indicates number of replications, line delineates deletion vs. duplication/copyback events, while color indicates count/10^6 mapped reads. Log coverage data shown above, with light blue indicating average coverage over three replicates and dark blue the standard deviation.

## DISCUSSION

The goal of these studies was to determine how and whether intracellular D-RNA populations differ from encapsidated populations, addressing whether D-RNAs can arise *de novo* without requiring packaging and subsequent accumulation through high MOI passaging. By sequencing early-passage, low MOI-derived stock, intracellular passage “1.5” virus, and subsequent encapsidated passage 2 virus, we have shown that D-RNA diversity and composition are fundamentally different between intracellular and resulting encapsidated RNAs, both *in vitro* and *in vivo*, for 5 alphavirus strains from 4 distinct species and 2 distinct phylogenetic groups. As a result of these studies, we additionally observed a novel conserved subtype of alphavirus D-RNA originating from recombination events in the subgenomic region, specifically between capsid/E3 and end of E1/3’UTR subgenomic regions, which is heavily enriched intracellularly but do not appear to be commonly packaged. We’ve called this new heterogeneous subpopulation subgenomic D-RNAs (sgD-RNAs).

Viruses, especially RNA viruses, are notorious for their ability to generate highly diverse populations, particularly in terms of point mutations that are generated during replication and may contribute to overall viral fitness within a host (62, 63). Although researchers tend to think of viral population dynamics in terms of encapsidated progeny viruses, there may be a role for intracellular RNA diversity that remains unexplored. Here, we show that D-RNA populations differ between intracellular and encapsidated factions in early viral passages. For all viruses tested, there were significantly more unique recombination events intracellularly than were packaged, indicating that only a portion of D-RNAs generated are packaged. Overall numbers of D-RNAs are additionally higher intracellularly, and thus Shannon’s entropy index *H* is significantly higher in intracellular samples, indicating greater alpha diversity intracellularly. These trends held for *in vivo* studies, where D-RNA populations from specific tissues were more diverse than those from serum. Broadly, this indicates a packaging bottleneck between intracellular and encapsidated populations, which likewise means that production of D-RNAs occurs *de novo* without a strict requirement for transmitted template. These results also distinguish packaged D-RNAs and D-RNAs produced *de novo* as two separate populations. Packaged D-RNAs have been shown to play multiple roles during viral infection, which range from interfering with replication of wild-type parental virus RNA and promoting viral persistence in tissue culture to stimulating the host immune response *in vivo*. However, unpackaged D-RNAs have not been appreciated as a distinct population before now and subsequently the role of these D-RNAs remains unclear. These studies therefore provide a critical foundation for further investigation of the role of *de novo* D-RNA production during viral infection by establishing this important distinction between packaged and unpackaged D-RNAs.

There is some evidence that recombination events leading to D-RNA formation are largely, though not exclusively, polymerase-driven (64–66) and are influenced by polymerase characteristics such as fidelity. For example, low-recombination variants of Senecavirus exhibited a high-fidelity polymerase phenotype (67), a low-fidelity SINV polymerase mutant showed increased rates of encapsidated D-RNA production (35), and a high-fidelity poliovirus polymerase mutant exhibited decreased rates of recombination (68). A popular model for recombination events leading to D-RNA formation is through a copy-choice mechanism, in which a polymerase retains the nascent RNA chain but switches templates during transcription (69, 70). In this model, it is generally thought that template-switching occurs during anti-sense strand synthesis, thus giving rise to a defective template prior to generation of daughter sense-strand RNA. This is a reasonable hypothesis for a few reasons, including: 1) anti-sense template is generally rare, affording few opportunities for recombination between discrete templates; and 2) to produce D-RNAs to transmissible levels, presumably a defective template would be required to generate a minimum copy number to increase packaging efficiency. Kirkegaard and Baltimore were also able to demonstrate this experimentally using a wild-type poliovirus and a guanidine-resistant poliovirus mutant, which showed preferential template-switching activity consistent with that predicted for the generation of recombinant negative-sense template RNA (69).

However, the data presented here suggests that template-switching may not occur exclusively during anti-sense template synthesis. Firstly, that alphavirus D-RNA production likely occurs *de novo* implies that transmitted template is not a strict requirement for D-RNA production. In addition, differential D-RNA diversity between intracellular and encapsidated alphavirus RNA was due almost entirely to recombination events in the subgenomic region encoding the structural proteins. The packaging signals of alphaviruses are thought to be in the nsP1 (71, 72) and, in the case of Semliki Forest virus, nsP2 genes (3) in the nonstructural gene cassette, thus the lack of packaging of these particular D-RNAs may be due to the fact that they are specifically on sgRNA transcripts and not genomic transcripts. This was confirmed by DRS, which showed that CHIKV sgDels were expressed primarily in sgRNA transcripts. Furthermore, since genomic and sgRNA transcripts are both transcribed from full-length negative-sense template RNA, then we would reasonably expect to see the same deletions in both genomic and subgenomic reads in long-read sequencing data if template-switching occurs during negative-sense strand synthesis; however, because no overlap in deletions was observed between genomic and subgenomic reads in either of our DRS data sets, we could not confirm the presence of a common defective template. Together, these results suggest that either: 1) template-switching can occur during either antisense- or sense-strand synthesis; or 2) sgD-RNAs are not generated through a copy-choice mechanism. Clustering of recombination junctions into “hotspots” supports the former.

Similar to previous work, we also observed substantial recombination activity in the 3’UTR of CHIKV. These CHIKV 3’UTR recombination events mostly consisted of duplications, especially in intracellular populations. Duplications in the CHIKV 3’UTR are well-documented and are thought to influence CHIKV host adaptability, particularly in the insect vector (36, 38, 73). Filomatori and Bardossy and colleagues additionally found that these recombination events arise from a mixture of homologous- and nonhomologous-template switching (39), although the relative quantities of each varied depending on host type. In addition to these, we have also shown that CHIKV, MAYV, SINV, and AURV all readily produce sgD-RNAs featuring recombination events in the subgenomic region, especially subgenomic deletions (sgDels) during early, low-MOI passages, as well as in a murine model of CHIKV infection. All sgD-RNAs share similar boundaries between all viruses tested, particularly deletions with the first junction occurring between capsid-E3 regions and the second occurring between E1-3’UTR regions. Because these populations arise regularly after infection not only with different CHIKV stock viruses but also during infection with heterologous alphaviruses, with unpredictable overlap between those observed among intracellular and encapsidated RNAs, this suggests they arise *de novo* and are consistently expressed at low levels intracellularly. The boundaries appear to be loosely conserved between the alphaviruses tested, which all either cause or are highly related to viruses that cause arthritogenic disease (56). Interestingly, 3’UTR events were relatively enriched in encapsidated factions compared to sgDels and other subgenomic recombination events, potentially indicating a different mechanism through which these events occur.

Intriguingly, although this study focused on arthritogenic alphaviruses, the general sgD-RNA boundaries correspond to a well-known recombination event between Eastern equine encephalitis virus (EEEV) and SINV that gave rise to the Western equine encephalitis virus (WEEV) species (56, 74). Little is known about the particulars of this recombination event, such as whether it occurred in an insect or mammalian host, but determining the mechanism of expression of sgD-RNAs among alphaviruses may help delineate the circumstances that gave rise to WEEV. This is especially important, as CHIKV is now co-circulating with MAYV (75), Una virus (75), and Eastern and Venezuelan equine encephalitis viruses since its introduction to the Western hemisphere in 2013, presenting new opportunities for heterologous alphavirus recombination events that may gave rise to novel recombinant alphavirus species.

A common concern regarding D-RNA research is that they may simply be an artifact of tissue culture, given the contrived passaging conditions from which most D-RNAs have been identified and characterized. Whether D-RNAs arise during natural infection, and what roles D-RNAs may play, is therefore an important question. RSV D-RNAs have been recovered from patient samples, and indeed have been correlated with an enhanced host-response to infection (17, 76), while similar findings have been described for IAV (19). For alphaviruses, it has previously been shown that salmonid alphavirus 3 forms many D-RNAs in wild salmon hosts (16), among which 6K deletion mutants are common (41). 6K deletion mutants have also been observed among VEEV RNA populations derived from sentinel hamster hosts (42). However, we have definitively shown that a pathogenic alphavirus, CHIKV, forms D-RNAs with substantial incapacitating deletions in a mammalian host. Importantly, many of the recombination events we observed *in vivo* were similar to ones observed *in vitro* during early passages, suggesting that expression of sgD-RNAs is a programmed phenomenon with potential biological functions, recapitulating observations made with IAV (21).

In all, the current study provides critical evidence of discrete types of D-RNAs, encapsidated/transmissible and intracellular/*de novo-*produced, as well evidence for a novel type of alphavirus D-RNA, sgD-RNAs. Further study is required to ascertain the origin of alphavirus sgD-RNAs and the factors leading to formative recombination events. There are two general hypothesis that are not mutually exclusive: 1) that sgD-RNAs are an artifact of another replicative process, such as the binding of host and/or viral proteins to viral RNA; and 2) that sgD-RNAs are a deliberate by-product of viral replication with specific cellular functions. Additionally, the function of alphavirus sgD-RNAs and other *de novo*-produced D-RNAs remains unknown, necessitating further investigation.

## ACKNOWLEDGMENTS

We would like to thank Scott Weaver for his support on this project, especially for the use of BSL-3 facilities and guidance for animal studies, as well as Dr. Kenneth Plante and the World Reference Center for Emerging Viruses and Arboviruses for providing additional viruses and input on best culture practices.

Funding for RML was provided by the Jeanne B. Kempner postdoctoral fellowship. A.R. and this work was supported by start-up funds from the University of Texas Medical Branch, Galveston.

## METHODS

### Tissue Culture

Vero cells (American Type Culture Collection, Manassas, VA) were maintained in Dulbecco’s modified Eagle medium (DMEM; Gibco) containing 1X Penicillin/Streptomycin/Amphocertin (PSA; Gibco, Waltham, MA) and 5% fetal bovine serum (FBS; Hyclone, Logan, UT). *Aedes albopictus* mosquito (C7/10) cells, kindly provided by Scott Weaver, were maintained in DMEM containing 1X PSA, 10% FBS, 1X non-essential amino acids (Sigma, St. Louis, MO), 1X sodium pyruvate (Gibco), and 5mL 1X tryptose phosphate buffer (Gibco).

### Viruses and infections

CHIKV 181/clone 25 (77), CHIKV 15561 (77), and MAYV IQU3056 (78) infectious clones were kindly provided by Dr. Scott Weaver. Low-passage AURV BeAr93375 (P2, C6/36) and SINV CGLT599 (P2, suckling mouse) isolates were provided by the World Reference Center for Emerging Viruses and Arboviruses (WRCEVA; Galveston, TX). Infectious clones were rescued as follows: first, plasmids were digested with appropriate endonuclease, and then digested plasmid was purified using phenol:chloroform:isoamyl alcohol. Then, 1ug digested plasmid was added to mMessage mMachine SP6 reaction (Ambion, Austin, TX) followed by DNA digestion and RNA clean-up using RNA Clean and Concentrator kit (Zymo, Irving, CA) following manufacturer’s instructions. Cleaned RNA was electroporated into Vero cells using a Neon electroporation device following manufacturer’s Vero protocol (Thermo Fisher Scientific, Waltham, MA). All viruses were amplified in C7/10 cells using MOI=0.1 to generate P1 stock virus; P0 and P1 stock titers were determined in vero or BHK21 cells using plaque assays as previously described (79).

Viral infections were performed as follows: virus was first diluted in serum/antibiotic-free DMEM to a target MOI of 2 infectious units/cell. Culture medium was removed from cells and virus-medium added for 1 hour, rocking plates every 15 minutes. Cells were then washed 3X with PBS, and normal culture medium added.

### RNA extractions

Intracellular RNA was collected 11-12 hours post-infection as follows: supernatant was removed from target wells, followed by 3x washes with 2 mL PBS, and then 500uL TRIzol (Invitrogen, Carlsbad, CA) was added directly to plate and incubated for 5 minutes prior to collection. Packaged RNA was collected 48 hours post-infection as follows: supernatant was collected, clarified by centrifugation for 5 minutes at 1000 rcf, and incubated in 7% PEG8000 NaCl overnight at 4°C. Precipitate was then pelleted by centrifugation at 3,200xg for 30 minutes at 4C and supernatant removed, pellet was resuspended in 100uL 10% FBS DMEM, and then incubated with 10ug RNase A for 2 hours at room temperature (to remove contaminating non-packaged RNAs). 500uL TRIzol was then added to RNase-treated sample and incubated for 5 minutes. For both intracellular and packaged RNAs, RNA was then extracted according to manufacturer’s instructions using Phasemaker tubes (Invitrogen). Additionally, ribosomal RNA was removed from intracellular RNA samples using Lexogen RiboCop kit according to manufacturer’s instructions (Lexogen Inc, Greenland, NH).

### Animal infections and RNA collections

Type 1 interferon-deficient IFNαR^-/-^ (A129) mice of varying ages and genders were kindly provided by Dr. Slobodan Paessler. Mice were infected with 10^3^ PFU CHIKV AF15561 in 10uL PBS in the left rear-footpad and monitored for weight and signs of disease. Animals were euthanized 4 days post-infection by CO2 overdose followed by cardiac puncture, whereupon blood, skeletal muscle from the infected and contralateral legs, heart, kidney, liver, and spleen: organs were placed in 300uL RNA later solution, while whole blood was centrifuged at 3,380 rcf for 5 minutes and serum collected and stored at −80 until further use. Upon processing, tissue was removed from RNAlater solution, rinsed in sterile PBS, and placed in a 2mL microcentrifuge tube with 300uL Buffer RLT and a stainless steel ball bearing. Samples were heat inactivated at 60C for 15 minutes, and then organs were homogenized using a TissueLyzer II (Qiagen, Hilden, Germany) shaking at 26 p/sec for 5 minutes. RNA was then extracted from organs in Buffer RLT using RNeasy Fibrous Tissue mini kits following manufacturer’s instructions (Qiagen), while RNA was collected from serum using TRIzol according to manufacturer’s instructions. Mouse studies were performed in an animal biosafety level 3 facility according to approved UTMB IACUC protocol #1708051. The University of Texas Medical Branch complies with NIH policy on animal welfare, the Animal Welfare Act, and all other applicable federal, state, and local laws.

### Next generation sequencing

ClickSeq libraries were generated as previously described (43, 44). Oxford Nanopore Technologies’ Direct RNA sequencing (DRS) libraries were constructed as follows: intracellular RNA was immediately ribo-depleted using Ribo-Zero kits (Illumina, San Diego, CA) upon RNA extraction and used within 24 hours of ribodepletion (stored on ice). DRS kits (Oxford Nanopore Technologies, Oxford, UK) were purchased directly from ONT and libraries were constructed using 250-300 ng of starting RNA according to the most recent version of the DRS protocols published by the manufacturer. The libraries were run on a MinION sequencer using R9.4.1 flow cells for 8-12 hours each.

### Bioinformatics

For AURV BeAr93375 and SINV CGLT599, corrected reference genomes were generated using *Pilon* based on publicly available sequences (80). For CHIKV 181/clone 25, CHIKV AF15561, and MAYV IQU3056 clones, the viral cDNA sequences provided by the Weaver lab were used for reference genomes. Reads were trimmed and quality filtered using the *fastp* tool’s default parameters (45), and then recombination events were identified and enumerated using our *ViReMa* (46) v1.5 integrated pipeline with the following inputs:

~~~
**[script location]**/ViReMa.py **[reference genome]**.txt **[input file]**.fastq
**[output file designation]**.sam --Output_Dir **[output folder]** --p **4** --
MaxIters **10** --MicroInDel_Length **5**
~~~

The resulting .txt files from *ViReMa* were used to calculate Shannon’s entropy (*H*), weighted by median nucleotide coverage, utilizing a previously described custom python script (44). Further, using *DESeq2* (47), for each virus recombination events were compiled, normalized, and expression changes evaluated based on *ViReMa* recombination output file. From these results, hierarchical clustering was performed using *Cluster* 3.0 (48), filtering for a minimum of 2 replicates with expression values >2; resulting trees were visualized using *TreeView* (49). Additionally, normalized data from DESeq2 were used to perform principle component analyses in *SigmaPlot*. ONT sequences were aligned to alphavirus genomes using *minimap2* (51)and read alignments extracted using a custom python script.

### Statistics

All statistics were performed in *SigmaPlot*. For Shannon’s entropy and unique/total D-RNA comparisons, data were tested for normality using the Shapiro Wilk test, followed by Student’s T-test comparing intracellular and encapsidated P2 samples. For animal data, data were tested for normality followed by one-way analysis of variance (ANOVA). Data formatted by *DESeq2* were used for principle component analyses performed in *SigmaPlot*.

### Data availability

All raw data files for Illumina and default quality-filtered/base-called data for direct RNA nanopore sequencing datasets associated with this manuscript are available on the SRA NCBI archive under study number PRJNA613616.

