## Supplemental figures for "Differential alphavirus defective RNA diversity between intracellular and encapsidated compartments is driven by subgenomic recombination events"

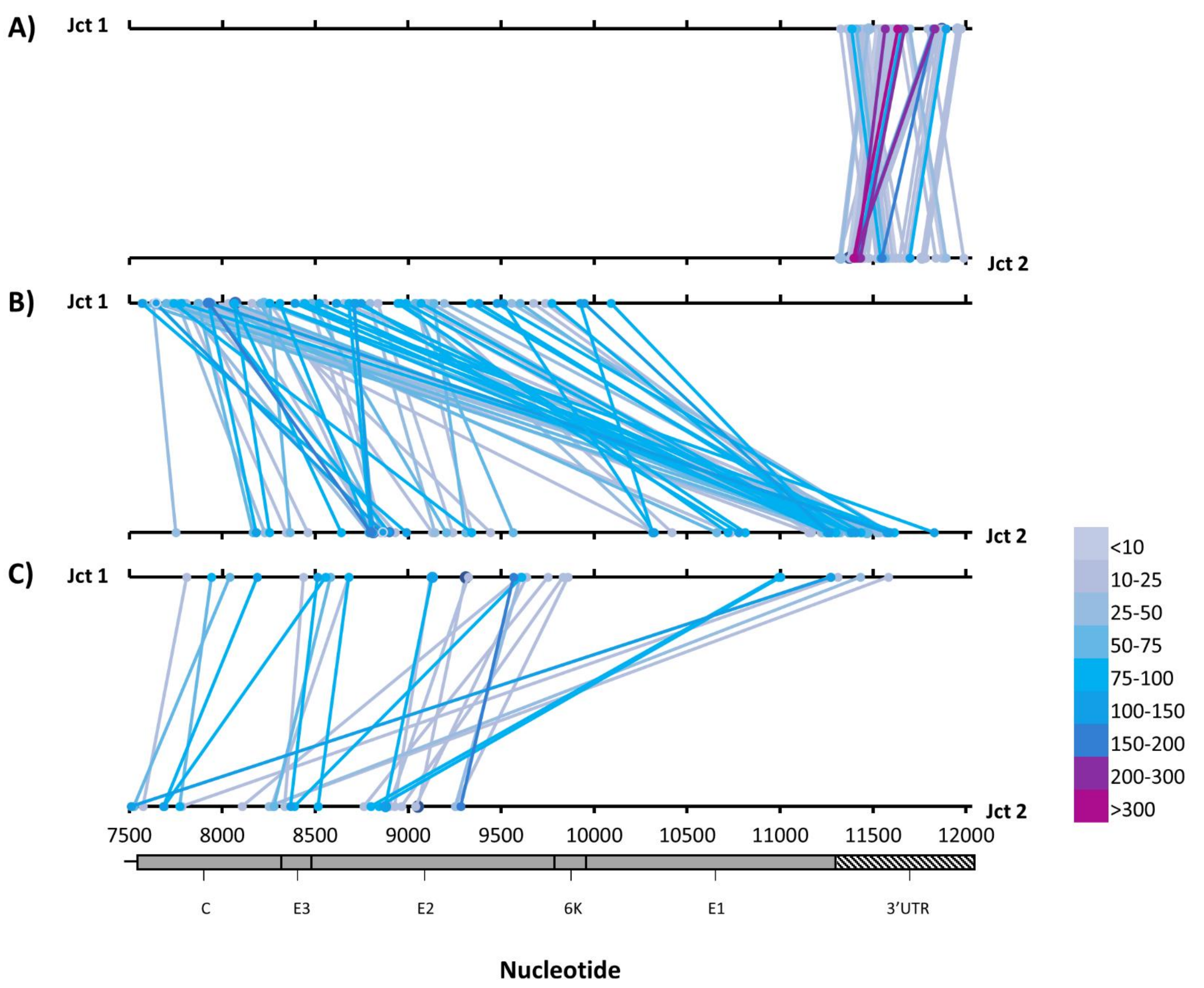


**Fig. S1 CHIKV subgenomic recombination junctions**. Three replicates of intracellular RNA from chikungunya virus (CHIKV)-infected Vero cells was collected 12 hours post-infection and then sequenced and evaluated for defective RNA expression using *ViReMa* v1.5. Recombinant count data were normalized to count per $10^6$ CHIKV-mapped reads, and then subgenomic recombination events (both junctions, Jct, occurring after nucleotide 7500) were averaged and then ranked from highest to lowest count; top 200 in total are shown. Top line represents donor site (Jct 1) and bottom line represents acceptor (Jct 2). **A)** 3'UTR-specific recombination events **B)** Subgenomic deletions **C)** Subgenomic insertion/duplication events.

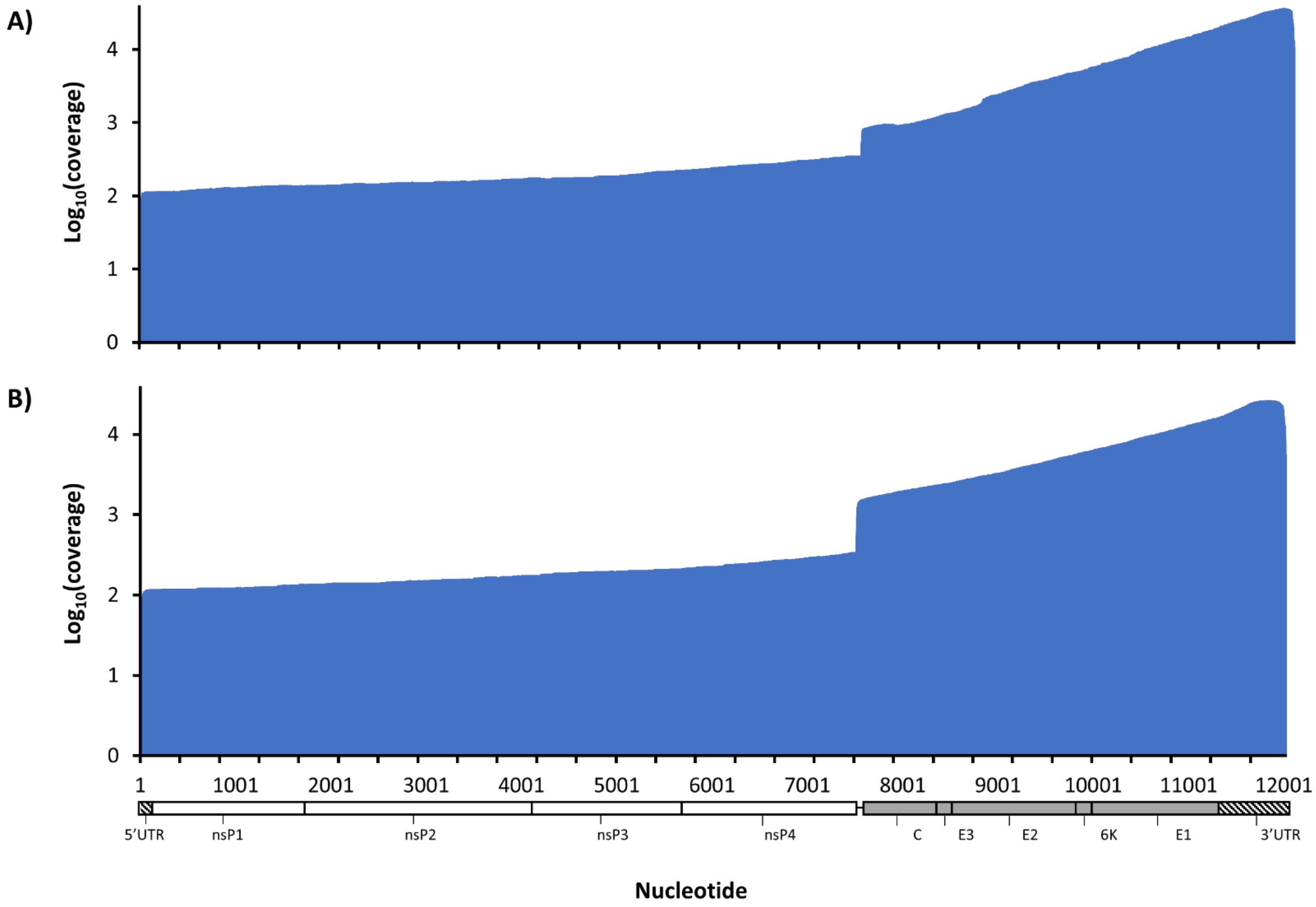


**Fig. S2 Coverage data for CHIKV MinION Direct-RNA Sequencing reactions.** Two replicates of intracellular RNA from chikungunya virus (CHIKV)-infected Vero cells was collected 12 hours post-infection and then two separate sequencing libraries were constructed using Oxford Nanopore Technologies' Direct RNA sequencing kit and sequenced on a MinION sequencer. After aligning reads to the CHIKV genome, $Log_{10}$ nucleotide coverage for both reactions (DRS1, **A**; DRS2, **B**) was plotted.

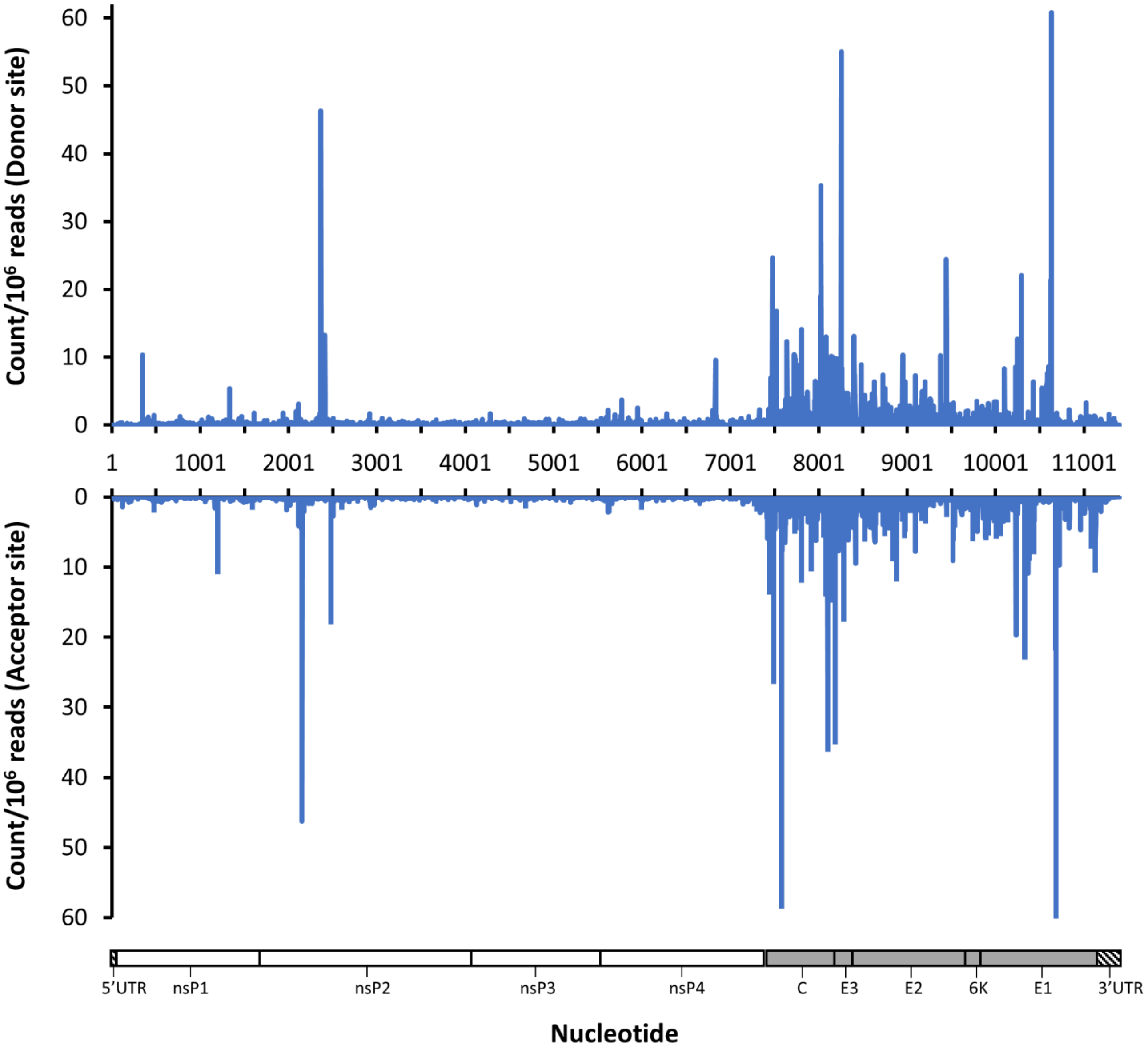


**Fig. S3 Recombination junction counts for MAYV D-RNAs**. Three replicates of intracellular RNA from Mayaro virus (MAYV)-infected Vero cells was collected 12 hours post-infection and then sequenced and evaluated for defective RNA expression using ViReMa v1.5. Recombinant count data were normalized to count per 106 MAYV-mapped reads, and then average nucleotide count for donor sites (Top) and acceptor sites (bottom) were plotted.

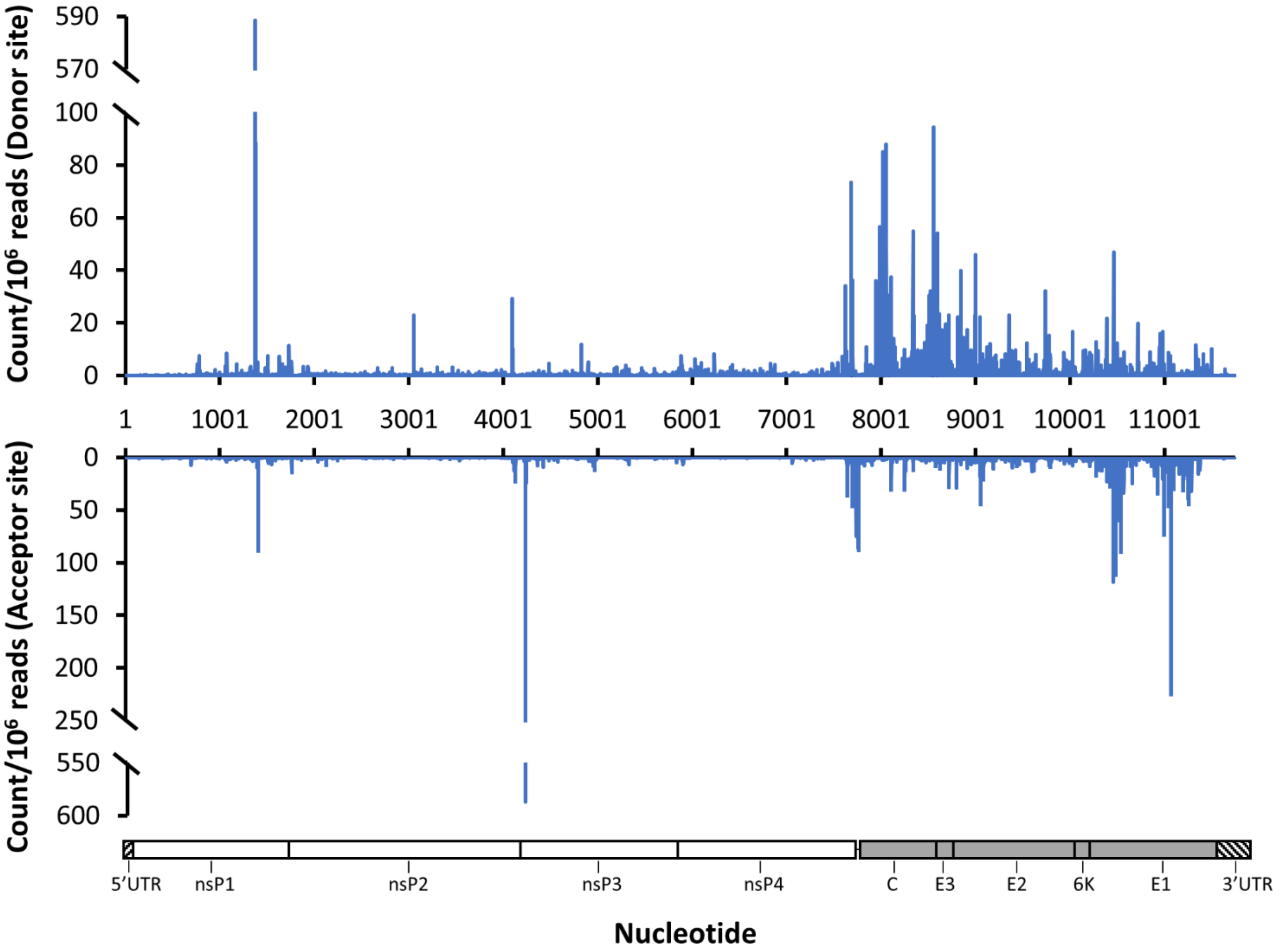


**Fig. S4 Recombination junction counts for SINV D-RNAs**. Three replicates of intracellular RNA from Sindbis virus (SINV)-infected Vero cells was collected 12 hours post-infection and then sequenced and evaluated for defective RNA expression using ViReMa v1.5. Recombinant count data were normalized to count per 106 SINV-mapped reads, and then average nucleotide count for donor sites (Top) and acceptor sites (bottom) were plotted.

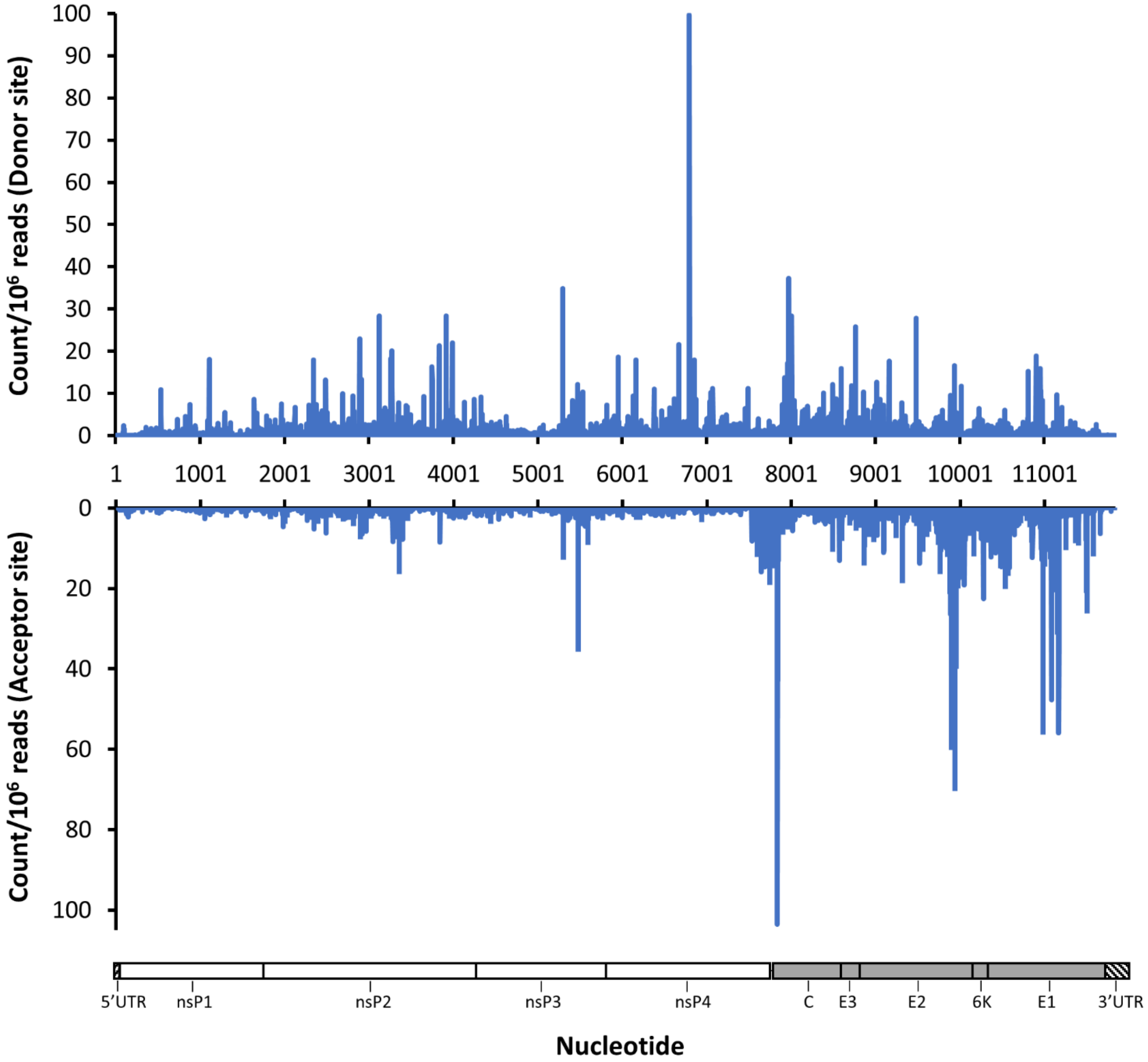


**Fig. S5 Recombination junction counts for AURV D-RNAs**. Three replicates of intracellular RNA from Aura virus (AURV)-infected Vero cells was collected 12 hours post-infection and then sequenced and evaluated for defective RNA expression using ViReMa v1.5. Recombinant count data were normalized to count per 106 AURV-mapped reads, and then average nucleotide count for donor sites (Top) and acceptor sites (bottom) were plotted.

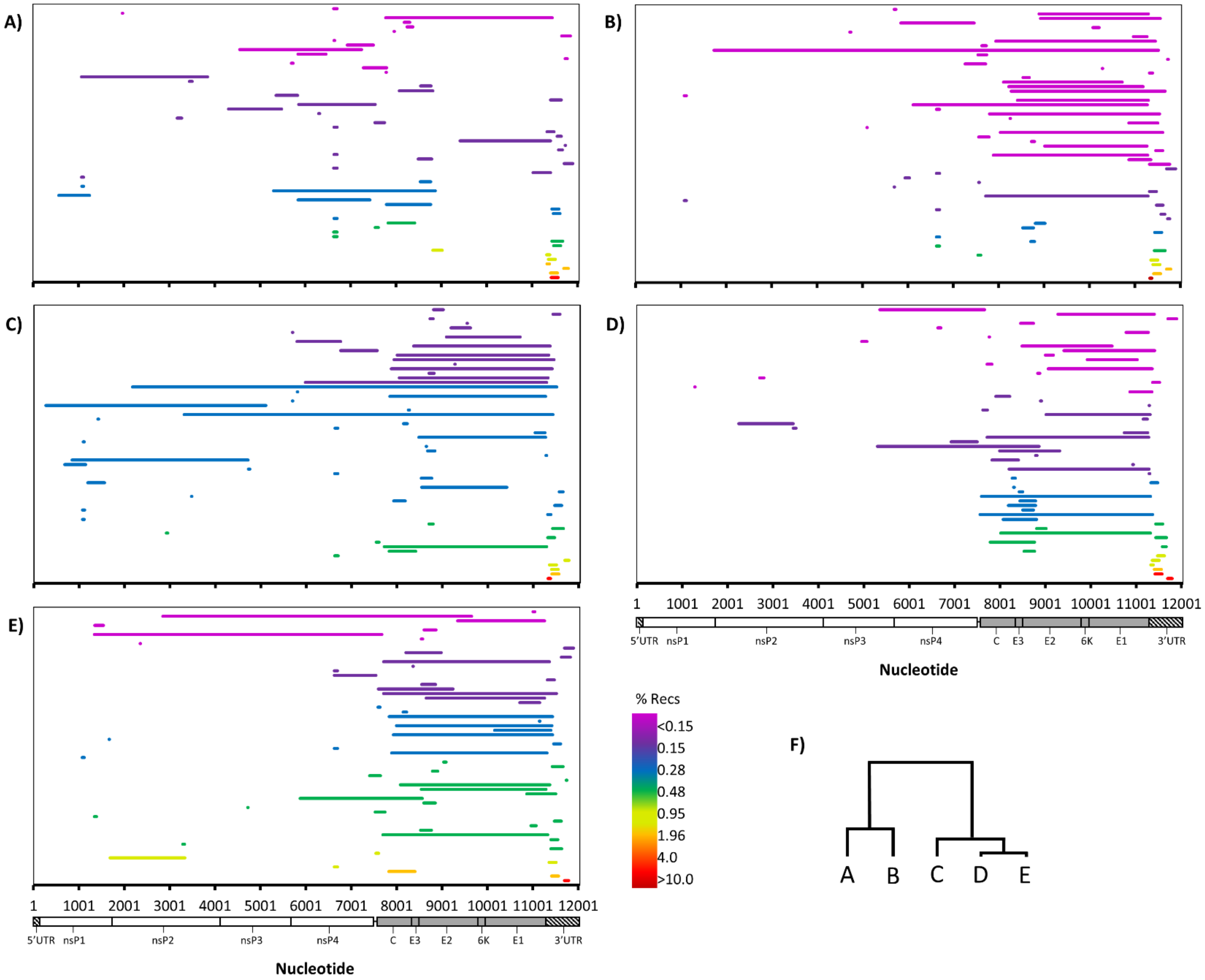


**Fig. S6. Top deletion events observed among CHIKV-infected mouse organs**. Four A129 mice were infected with CHIKV AF15561 in the left rear footpad, and then animals were humanely euthanized and serum and organs collected 4 days post-infection. RNA was then extracted from serum and tissues, and then sequenced on an Illumina NextSeq550. Defective-RNAs were identified and quantified using *ViReMa v1.5.* Top 60 deletions for Mouse 2 serum **(A)**, contralateral leg muscle **(B)**, injection site leg muscle **(C)**, cardiac muscle **(D)**, and mouse 1 kidney **(E)** (a low overall number of recombination events were observed in M2-kidney sequencing reaction, thus to compare deletions M1 kidney was used) are shown. **F)** Hierarchical analysis of D-RNA expression and organ clustering for mouse 1.

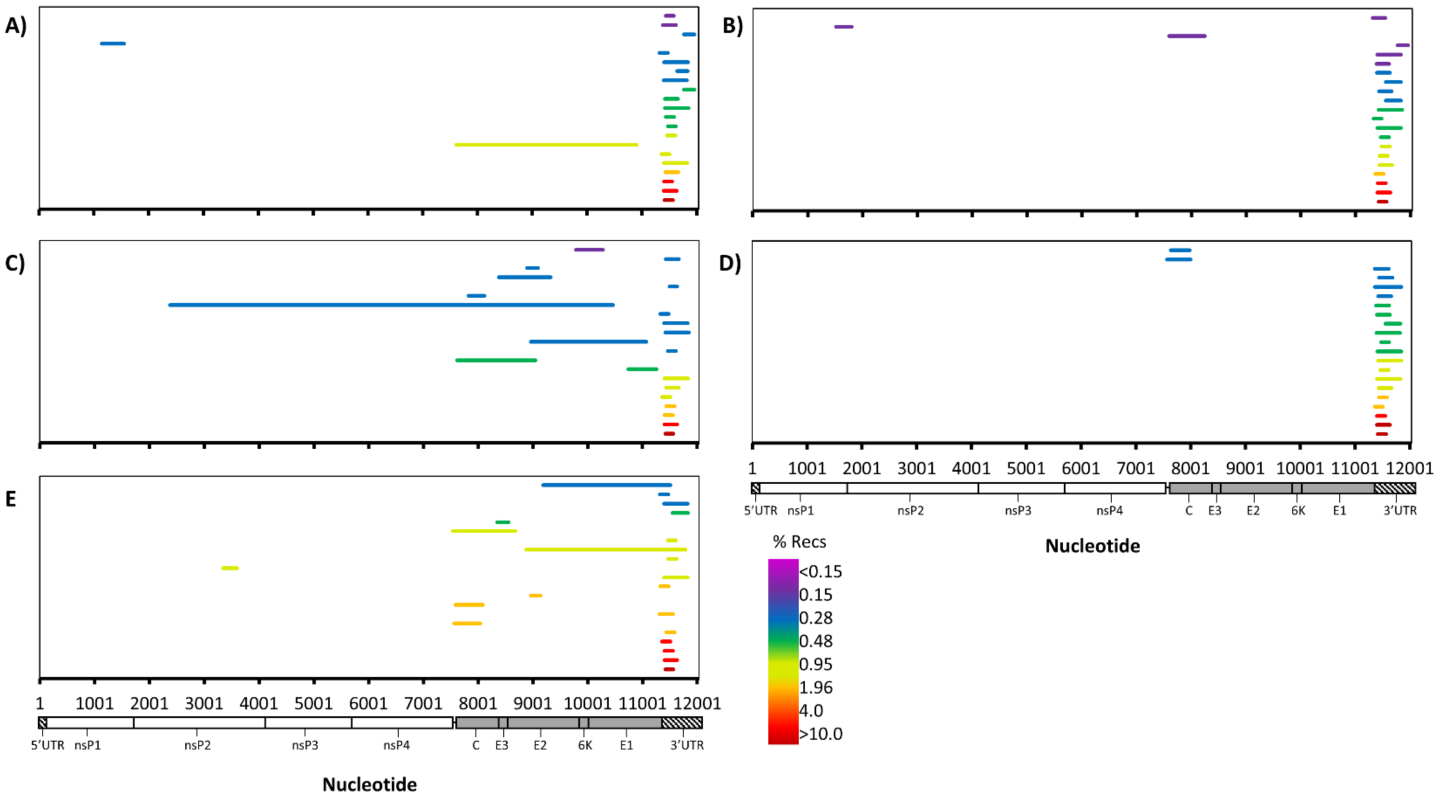


**Fig. S7.. Top insertion/duplication events observed among CHIKV-infected mouse organs.** Four A129 mice were infected with CHIKV AF15561 in the left rear footpad, and then animals were humanely euthanized and serum and organs collected 4 days post-infection. RNA was then extracted from serum and tissues, and then sequenced on an Illumina NextSeq550. Defective-RNAs were identified and quantified using *ViReMa v1.5.* Top 21 insertion/duplication events for Mouse 2 serum **(A)**, contralateral leg muscle **(B)**, injection site leg muscle **(C)**, cardiac muscle **(D)**, and kidney **(E)** are shown
